# Plasmacytoid dendritic cells are dispensable or detrimental in murine systemic or respiratory viral infections

**DOI:** 10.1101/2024.05.20.594961

**Authors:** Clemence Ngo, Camille Pierini-Malosse, Khalissa Rahmani, Michael Valente, Nils Collinet, Gilles Bessou, Capucine Guerry, Manon Fabregue, Solene Mathieu, Sarah Sharkaoui, Sophie Mazzoli, Amandine Sansoni, Frederic Fiore, Caroline Laprie, Lena Alexopoulou, Mauro Gaya, Claude Gregoire, Achille Broggi, Sarah Wurbel, Réjane Rua, Narjess Haidar, Pierre Milpied, Bertrand Escalière, Thien Phong Vu Manh, Mathieu Fallet, Lionel Chasson, Hien Tran, Thomas Baranek, Marc Le Bert, Bernard Malissen, Ana Zarubica, Marc Dalod, Elena Tomasello

**Affiliations:** Aix-Marseille University, CNRS, INSERM, CIML, Centre d’Immunologie de Marseille-Luminy, Turing Center for Living Systems, Marseille, France; Centre d’Immunophénomique, Aix Marseille Université, Inserm, CNRS, PHENOMIN, CELPHEDIA, Marseille, France; Laboratoire Vet-Histo, 122 Av Joseph Vidal, 13008 Marseille; Department of Microbiology, Immunology and Glycobiology, Institute of Laboratory Medicine, Lund University, 223 62 Lund, Sweden; Centre d’Étude des Pathologies Respiratoires (CEPR), Institut National de la Santé et de la Recherche Médicale (INSERM) Unité Mixte de Recherche 1100, Faculté de Médecine, Université de Tours, Tours, France; Immuno-Neuro Modulation, Université d’Orléans Campus CNRS, 3B rue de la Ferollerie, 45071 ORLEANS, France

## Abstract

Plasmacytoid dendritic cells (pDCs) are major producers of type I/III interferons. As interferons are crucial for antiviral defense, pDCs are assumed to play an essential role in this process. However, robust evidence supporting this dogma is scarce. Genetic or pharmacological manipulations that eliminate pDC or disrupt their interferon production often affect other cells, confounding interpretation. To overcome this issue, we engineered pDC-less mice that are specifically and constitutively devoid of pDCs by expressing diphtheria toxin under coordinated control of the *Siglech* and *Pacsin1* genes, uniquely co-expressed in pDCs. pDC-less mice mounted protective immunity against systemic infection with mouse Cytomegalovirus and showed higher survival and less lung immunopathology to intranasal infection with influenza virus and SARS-CoV2. Thus, contrary to the prevailing dogma, we revealed that pDCs and their interferons are dispensable or deleterious during several viral infections. pDC-less mice will enable rigorously reassessing the roles of pDCs in health and disease.

## Introduction

Vertebrate antiviral immunity heavily depends on type I and type III interferons (IFN-I/IIIs)^1^. IFN-I/IIIs promote all the three arms of antiviral immune defenses: intrinsic, innate and adaptive immunity. IFN-I/III binding to their respective receptors activate a major common pathway involving the activation by phosphorylation of three transcription factors, STAT1, STAT2 and IRF9, and their hetero-trimerization into the complex ISGF3 (interferon stimulated gene factor 3). ISGF3 binds to IFN-stimulated response elements (ISRE) in genomic promoter or enhanceantiviral intrinsic immunity in putatively all nucleated cells of the body^1^. IFN-Is also contribute to the activation or functional polarization of immune cells, hence promoting protective innate and adaptive anti-viral immune responses^1^.

IFN-I/III responses are generally beneficial during viral infections. However, when deregulated, these responses can turn deleterious^1^. This occurs when IFN-I/III are inappropriately induced in certain inflammatory or autoimmune diseases, in the absence of detectable viral infection. It can also happen during certain viral infections, if their induction is not properly controlled in time and space, as their proinflammatory effects can promote virus-induced immunopathology, contribute to immunosuppression, or increase susceptibility to secondary bacterial infections by preventing the induction of proper adaptive immune responses or delaying the healing of infection-induced tissue damage at epithelial barriers^1, 2, 3^. One strategy to promote the beneficial effects of IFN-I/IIIs and limit their harmful consequences would be to determine whether and how these distinct functions are linked to the modalities of their production. This includes identifying which cell types produce IFN-I/IIIs, how, when, where and with what functional consequences. Indeed, if specific cell types or sensing pathways are driving deleterious IFN-I/III responses in certain viral infections or autoimmune/inflammatory diseases, their specific targeting would allow dampening detrimental inflammation or misfired immune responses while preserving protective antiviral immunity. This approach would improve disease treatment by avoiding the general immunosuppression and the associated risk of infection caused by broad anti-inflammatory drugs such as corticosteroids or JAK/STAT inhibitors.

Amongst cellular sources of IFN-I/IIIs, plasmacytoid dendritic cells (pDCs) are special because they rapidly produce very high amounts of all subtypes of IFN-I/IIIs upon engulfment and endosomal sensing of nucleic acids derived from viral particles or material from infected cells, while being themselves highly resistant to viral infection^1, 4^. Specifically, pDCs express the endosomal TLR7 and TLR9 receptors that are able to recognize viral single stranded RNA and unmethylated CpG DNA sequences, respectively. TLR7/9 engagement induces IFN-I/III production via a MYD88-IRAK4-IRF7 signaling pathway^1^. In contrast, infected cells sense cytosolic or nuclear genome intermediates from the viruses replicating endogenously in their cytosol or nucleus, through specific helicases or the cGAS enzyme converging on the activation of a STING-IRF3 signaling pathway promoting the production of IFN-β and only few subtypes and low amounts of IFN-αs, with susceptibility to inhibitory viral immune-evasion mechanisms ^1, 5^. Hence, it is currently thought that pDCs generally play a critical and beneficial role in antiviral immunity as a major source of IFN-I/IIIs early after infection, indispensable for the reinforcement of intrinsic antiviral immunity and for proper orchestration of innate and adaptive immune responses^5, 6, 7, 8^. Yet, as we reviewed recently^4^ and will summarize in the next paragraph, direct and robust experimental evidence supporting this dogma is scarce, in a large part because methodological bottlenecks are preventing rigorous and definitive evaluation of the specific role of pDCs in antiviral immunity both in humans and in mice.

In humans, primary immune deficiencies abrogating pDC ability to produce IFN-I/IIIs, including loss-of-function mutations in TLR7, MYD88, IRAK4 or IRF7, have been associated primarily with a heightened susceptibility to *Mycobacterium tuberculosis* or a few other pyogenic bacteria but not to viral infections, except with the respiratory viruses Influenza and SARS-Cov2^9, 10, 11, 12, 13, 14, 15, 16, 17^. This contrasts with the much broader susceptibility to viral infections of patients suffering from inborn errors in genes necessary for IFN-I or IFN-III responses, such *as STAT1, STAT2, IRF9, IFNAR1* or *IFNAR2*^18, 19^. Hence, these data show that pDCs but not IFN-I/III responses are largely redundant for antiviral immunity in modern humans in the current hygiene and health care context. In mice, contrasting results were obtained regarding the consequences on antiviral immunity of the genetic inactivation of *Tlr7/9* or *Myd88,* depending on the combination of the virus and mouse strains studied, on differences in the inoculum dose or route, as well as on the type of readout measured. Indeed, the analysis of intrinsic antiviral immunity and side-by-side comparisons with mice deficient for IFN-I/III responses were seldom performed^1, 20^. In any case, whether in humans or mice, genetic inactivation of TLR7/9, MyD88, IRAK4 or IRF7 affects other cells and biological processes beyond pDC-derived IFN-I/III production. For example, TLR7 and IRF7 are also expressed and functional for IFN-I induction in monocytes and macrophages, and Myd88 is required for the responses to IL-1 cytokine family members in many cell types^4^.

Several approaches have been developed to deplete pDCs *in vivo* in mice to assess the functional impact of their loss. However, most of these approaches are not specific enough and have off-target effects. This confounds the rigorous interpretation of the observed phenotypes, making it difficult to determine whether they are specifically attributable to pDCs or to other cell types that are also directly impacted by the mutation or treatment, as summarized in a recent review^21^. For example, the cell types that are directly affected beyond pDCs include, many immune lineages in *Ikaros^L/L^* mice^22^; DC precursors and subpopulations of DCs and macrophages in *Siglech*-hDTR mice^23, 24^; subpopulations of DCs, macrophages and possibly B cells in *Itgax^Cre^*;*Tcf4^-/fl^*mice^25, 26, 27, 28, 29^, plasma cells, subpopulations of activated B cells, DCs, monocytes and macrophages in mice treated with anti-BST2/PDCA1 depleting antibodies. Studies performed using these tools supported the notion that pDCs were a major source of IFN-Is in infected mice, but varied in the assessment of the effective contribution of pDCs to anti-viral immunity, from beneficial to dispensable or even detrimental, depending on the infection route and the combination of virus and mouse strains^4^. Currently, pDC depletion can be achieved with high specificity and efficacy in one mutant mouse model, the hBDCA2-DTR mice, upon diphtheria toxin (DT) administration^30^. However, in these mice, the triggering of hDTR on pDCs upon diphteria toxin administration induces their production of IFN-I before causing cell death^31^. This might confound the interpretation of the requirement of pDCs in antiviral immunity or IFN-I responses when using this model. Moreover, repeated DT injections in hBDCA2-DTR mice have been reported to induce a severe morbidity proposed to be dependent on chronic pDC IFN-I production^31^. Therefore, there is still an unmet scientific need to determine rigorously whether and how pDCs modulate host antiviral defense *in vivo,* by selectively depleting pDCs while limiting off target and confounding effects.

Here, we reported the generation and characterization of pDC-less mice, a new mutant mouse model allowing specific and constitutive depletion of pDCs. pDC-less mice were engineered through an intersectional genetic strategy targeting diphtheria toxin A (DTA) expression in cells co-expressing the genes *Siglech* and *Pacsin1*, which we had previously validated as pDC-specific with a fluorescent reporter mouse model^32^. pDC-less mice were confirmed to be specifically devoid of pDCs in all tissues examined both at steady state and during viral infections. Yet, pDC-less mice remained able to establish protective anti-viral immunity during systemic infection with Mouse Cytomegalovirus (MCMV), a natural rodent pathogen. Moreover, pDC-less mice showed an enhanced resistance to respiratory infections with the influenza virus strain Scotland H3N2^33^ or with SARS-CoV2 that induce a severe lung immunopathology associated to a local cytokine storm. Moreover, IFN-I production by pDCs was deleterious in the Influenza infection model. Thus, our study revealed a detrimental role of pDCs and their IFN-I production during respiratory viral infections. In summary, we report here the generation of a novel mutant mouse model specifically and constitutively devoid of pDCs and its use demonstrating that pDCs are either dispensable or detrimental in murine systemic or respiratory viral infections. This challenges the prevailing dogma whereby pDCs generally play a crucial protective role in antiviral immunity. This new mouse model will be a critical tool for future studies aiming at rigorously determining the physiological functions of pDCs and their molecular regulation, in other models of infections and in inflammatory or autoimmune diseases.

## Results

### The pDC-less mice are constitutively and selectively devoid of pDCs

We recently generated *Siglech*^iCre^ x *Pacsin1*^LSLtdTomato^ double knock-in mice, called pDC-Tom, in which tdTomato fluorescent protein was exclusively expressed in “bona fide” pDCs^32^. Specific pDC targeting was achieved using intersectional genetic approach based on the co-expression of *Siglech* and *Pacsin1* genes uniquely in mouse pDCs. We adapted our two gene-based strategy by generating *Pacsin1*^LoxP-STOP-LoxP-DTA^ (*Pacsin1*^LSL-DTA^) knock-in mice, in which a floxed cassette containing the gene encoding the A subunit of the diphtheria toxin (DTA) was located at the 3’ of *Pacsin1* gene (**Fig. 1a**). *Pacsin1*^LSL-DTA^ were then bred with *Siglech*^iCre^, thus allowing the removal of STOP sequence and lethal expression of DTA exclusively in Siglech^+^Pacsin1^+^ cells, corresponding to pDCs. Indeed, pDCs were ablated in both lymphoid and non-lymphoid organs of double heterozygous *Siglech*^iCre^; *Pacsin1*^LSL-DTA^ mice, hereafter called pDC-less mice (**Fig. 1b** and **Extended Data Fig. 1a**). pDC-less mice developed and grew normally **(Extended Data Fig. 1b)**. Moreover, the cell numbers isolated from lymphoid and non-lymphoid organs of control versus pDC-less mice were comparable (**Fig. 1c**). The development of several other lymphoid and myeloid lineages was not affected, as exemplified in the spleen (**Fig. 1d-e**), thus confirming the selective ablation of pDCs. As pDCs and tDCs share a common hematopoietic precursor ^29^, we extended our analysis by using an optimized staining panel allowing to discriminate “bona fide” pDCs from other cDCs and tDC^hi^ and tDC^lo^ cell subsets expressing high or low levels of CD11c, respectively^27, 29^ (**Extended Data Fig. 1c).** We confirmed the lack of “bona fide” CD11c^lo^ Bst2^hi^ Ly6D^+^ CX3CR1^neg^ pDCs^32^ in our pDC-less mice (**Fig. 1f**). cDC2s were unaffected, while cDC1s were significantly increased in pDC-less mice, consistent with the previously reported enhanced expansion of cDC1s when the development of pDC-committed precursors is inhibited^34^. Finally, tDC^lo^/pDC-like cells were slightly, but significantly reduced, consistent with low, but detectable Tomato expression in late pDC-committed precursors isolated from pDC-Tom mice^32^, likely including recently described pro-pDC/tDC precursors^29, 32^. Altogether, our results demonstrate that, in pDC-less mice, pDCs are selectively and constitutively absent.

**Fig. 1:**
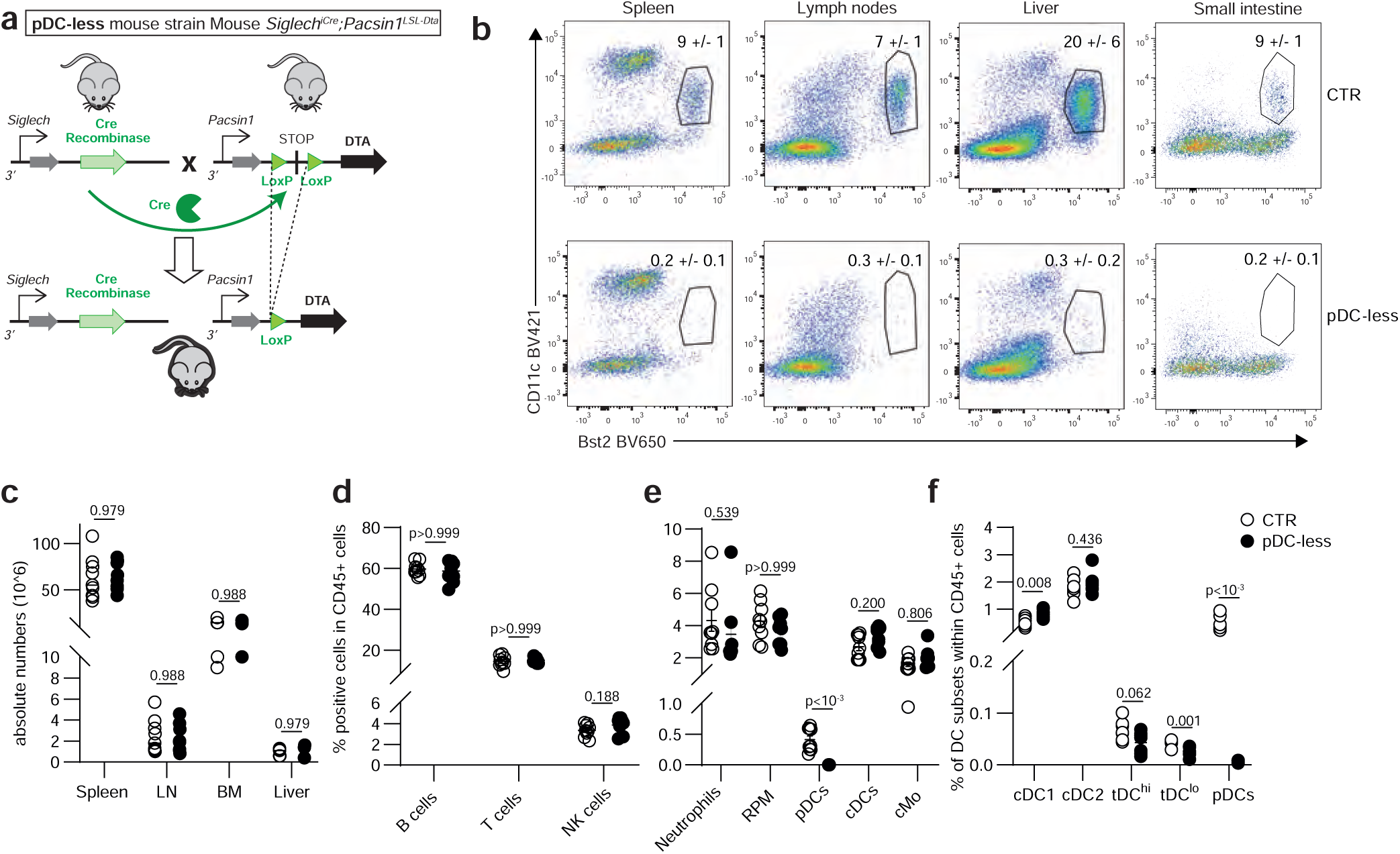
The pDC-less mice allow selective and constitutive depletion of pDCs in different organs. a, Scheme illustrating the strategy used to generate pDC-less mice. LoxP is the sequence recognized by Cre recombinase. ‘STOP’ corresponds to a transcriptional stop sequence. DTA= Diphtheria Toxin Subunit A. **b,** Dot plots showing the absence of lin-CD11b-CD11c^int^ Bst2^hi^ pDCs in cells isolated from indicated organs of control (CTR) vs pDC-less mice. pDC percentages (mean +/- s.e.m.) were pooled from two independent experiments (n=5 for each strain). **c,** Absolute numbers of total cells isolated from the indicated organs of CTR vs pDC-less mice (n= 9 for the spleen and lymph nodes (LN), n=5 for the bone marrow (BM) and liver). **d-f**, Percentages within CD45+ splenic cells of indicated lymphoid (**d**) and myeloid (**e,f**) lineages isolated from control vs pDC-less mice. RPM: red pulp macrophage cMo: conventional monocyte. The data shown (mean + s.e.m.) are pooled from two independent experiments (n = 9). An unpaired and nonparametric multiple t-test (Mann-Whitney) with Holm-Sidak method for correction was used for the statistical analysis.

### During MCMV infection, pDCs are the main source of IFN-I, but they are dispensable for cell-intrinsic and innate anti-viral immunity

During systemic infection by mouse cytomegalovirus (MCMV), a natural rodent pathogen, IFN-Is are mainly produced by pDCs^30, 35, 36, 37^. Indeed, both IFN-α and IFN-β were undetectable in the sera of MCMV-infected pDC-less mice at the peak of the IFN-I production detectable in infected control mice (1.5 days after infection). However, the production of other pro-inflammatory cytokines, such as IL-12, TNF, IFN-γ, CXCL10 and CCL2 was unaffected, thus supporting the existence of other sources of these cytokines redundant with pDCs (**Fig. 2a** and **Extended Data Fig. 2a**). We then evaluated the impact of the loss of pDCs and, especially of their IFNs, on anti-viral immunity. pDC-less mice survived MCMV infection, even at high viral loads (**Fig. 2b** and **Extended Data Figure 2b**) and efficiently controlled viral dissemination in all organs analyzed (**Fig. 2c**). Next, we analyzed the kinetics of ISG expression during MCMV infection. In infected control mice, the *Mx2* gene was induced at higher levels in the spleen than in the lungs and in the liver; its expression peaked in the spleen and the lungs at 1.5 days after infection, while it was delayed of 12 hours in the liver (**Extended Data Figure 2c**). When compared to uninfected mice, the expression of the ISGs *Mx2*, *Ifit2*, *Irf7* and *Isg15* was upregulated with similar kinetics in the organs of MCMV-infected control and pDC-less mice (**Figure 2d** and **Extended data Figure 2d**). However, we observed a high variability for each genotype and organ, with a trend for pDC-dependent expression of *Irf7* in the spleen and of *Mx2* and *Isg15* in the liver. In contrast, in infected *Ifnar1*^KO^ mice, the expression of these ISGs was downregulated in most of the organs studied. *Il28b* expression was reduced in a pDC-dependent manner in the spleen, but not in the liver of MCMV-infected mice (**Extended Data Figure 2d**), consistent with enhanced expression of *Il28a* and *Il28b* genes in IFN-I-producing splenic pDCs^37^. IFNs produced by pDCs during MCMV infection promote the activation of NK cells and cDCs, two innate immune lineages crucial for the induction of efficient responses against this virus^20, 30, 38, 39, 40^. However, granzyme B, IFN-γ and CD69 were similarly upregulated in splenic NK cells isolated from MCMV-infected control and pDC-less mice, regardless of the time point analyzed after infection (**Figure 2e**). Moreover, splenic cDC1s and cDC2s isolated from MCMV-infected control and pDC-less mice expressed comparable amounts of the IFN-induced CD86 activation marker (**Extended Data Fig. 2e**).

**Fig. 2:**
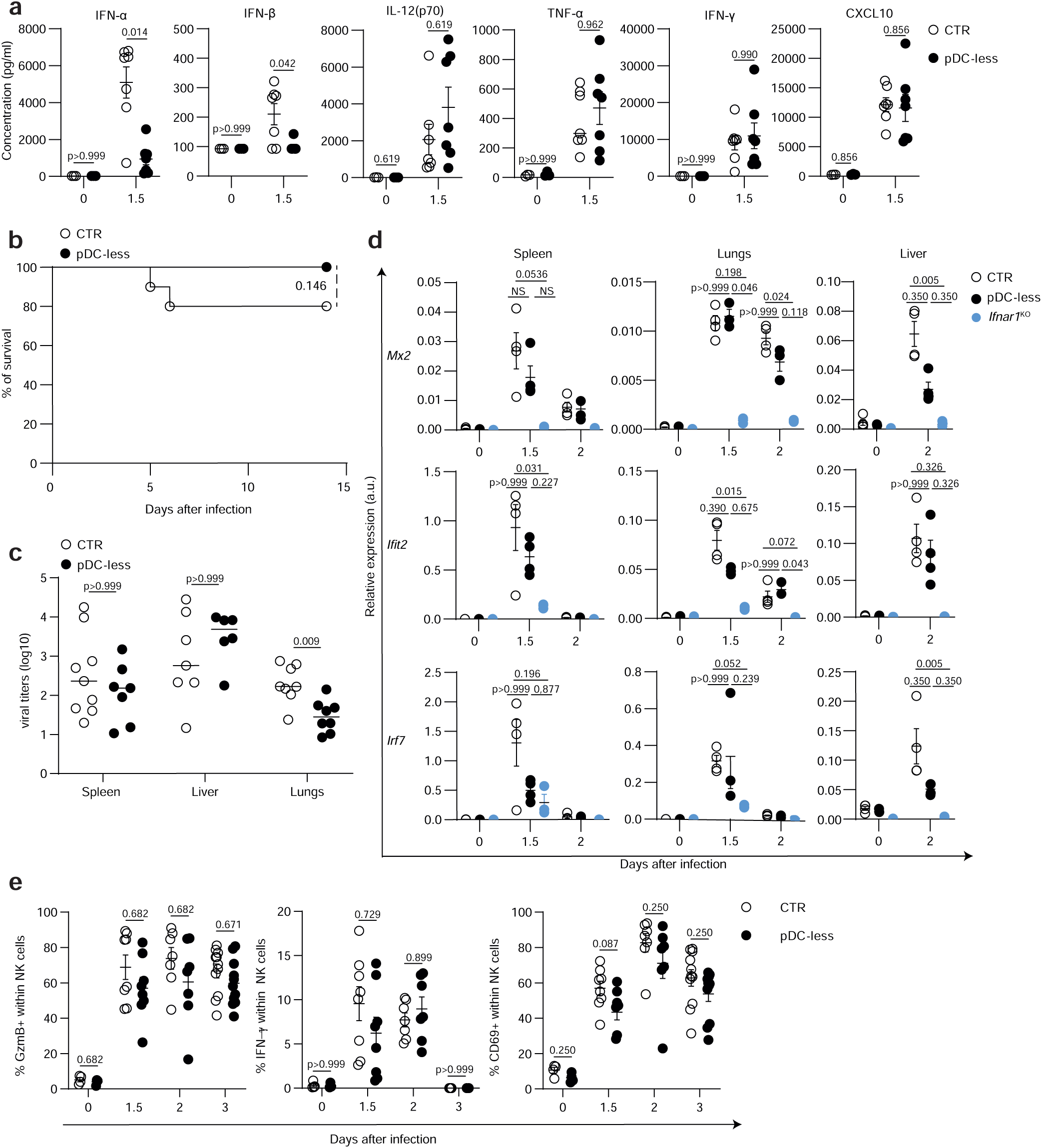
pDCs exert a dispensable role during MCMV infection. Control and pDC-less mice were infected (i.p) with 1x10^5^ PFU of MCMV Smith and analyzed at indicated time points after infection. **a,** Cytometric bead arrays was performed to quantify cytokine concentrations in the sera isolated from uninfected (0) and 36h infected (1.5) control and pDC-less mice. Each dot represents an individual animal. The data shown (mean +/- s.e.m.) are pooled from two independent experiments (n=3 for 0, n=7 for 1.5). An unpaired and nonparametric multiple t-test (Mann-Whitney) with Holm-Sidak correction method was used for the statistical analysis. **b,** Control and pDC-less mice were infected with 1.4 x10^5^ PFU of MCMV Smith and daily monitored for their survival, as shown in Kaplan-Meier curve (n= 10 for control, =8 for pDC-less). Log-rank (Mantel-Cox) test was used for statistical analysis. **c,** Viral titers were measured by RT-qPCR in the spleen, liver and lungs isolated from control and pDC-less mice 3 days after MCMV infection. Data shown are pooled from two independent experiments (9 CTR and 7 pDC-less for the spleen, 7 CTR and 6 pDC-less for the liver, n=8 for both strains for the lungs). Means are indicated by a horizontal bar. An unpaired and nonparametric multiple t-test (Mann-Whitney) with Holm-Sidak correction method was used for the statistical analysis. **d,** Expression levels of indicated genes was analyzed by RT-qPCR in indicated organs isolated from uninfected or MCMV-infected CTR, pDC-less and *Ifnar1*^KO^ mice at indicated days after infection. Data were normalized to expression level of *Actin* gene. The data (mean + s.e.m.) shown are from two independent experiments (n=4 for each condition at each time points). A nonparametric One-Way ANOVA (Kruskal-Wallis test with Dunn’s correction) was used for the statistical analysis. **e,** The percentages of GranzymeB (GzmB)^+^, IFN-γ^+^ and CD69^+^ cells within splenic NK cells (mean +/- s.e.m.) were determined in control and pDC-less mice at indicated days after MCMV infection (n=4 for 0, =8 for 1.5, =7 for 2 and =10 for 3 days after infection for both strains of mice). An unpaired and nonparametric multiple t-test (Mann-Whitney) with Holm-Sidak correction method was used for the statistical analysis.

Altogether, our results thus demonstrate that, during systemic MCMV infection, pDCs are dispensable for mounting protective intrinsic and innate anti-viral immunity, despite being the main IFN-I source.

### pDCs and the response to IFN-I are both detrimental in respiratory infection by influenza A virus

pDCs of patients bearing loss-of-function alleles of IRF7 were defective in IFN-I production in vitro upon TLR7 triggering and thus considered critical for the enhanced susceptibility of these patients to the respiratory influenza A virus (IAV) infection^12, 14^. In IAV-infected pDC-depleted mice, the scenario appeared more complex, supporting evidence of dispensable, protective or even detrimental functions of pDCs^41, 42, 43, 44, 45, 46^. However, in all mouse studies, other cells, besides pDCs, were depleted, thus confounding result interpretation. Moreover, both mouse genetic background and viral strain differentially affected mouse resistance to IAV infection. Contrary to 129 mice, C57BL/6 mice generally did not develop IFN-I-dependent immunopathology upon respiratory infections with most IAV strains, likely due to their relatively low production of IFN-I^44^. Thus, to address the potential pathogenic role of pDCs and their IFN-I production using our pDC-less mice, we used the IAV strain (H3N2 A/Scotland/20/74) known to induce a severe, Myd88-dependent, pneumonia in C57BL/6 mice^33^. When infected with the LD50 of this strain, half of the control mice died, whereas all pDC-less mice survived (**Fig. 3a** and **Extended Data Fig. 3a**). We extended our analyses to hBDCA2-DTR mice^30^. Upon DT administration, most of the IAV-infected hBDCA2-DTR mice survived, whereas 80% of the control C57BL/6 mice died (**Fig. 3b** and **Extended Data Fig. 3b**). Thus, in two distinct mutant mouse strains allowing their selective depletion, pDCs were deleterious during respiratory infection with IAV A/Scotland/20/74. IAV replication in the lungs did not differ between control and pDC-less mice, as observed by analyzing the expression of IAV *M1* gene at different time points after infection (**Fig. 3c**). IAV was undetectable in the brain of infected mice and viral titers were not modified in the absence of pDCs (**Supplementary Fig.1a**). The expression of the ISGs *Irf7*, *Isg15* and *Rsad2* was similarly induced in the lungs and, at lower extent, in the brains of infected control and pDC-less mice (**Fig. 3d-e and Supplementary Fig.1b**). Moreover, in the brains, we detected no significant difference in the frequency of mature (NeuN+) and activated (c-Fos+) neurons, microglia (Iba1+) and astrocytes (GFAP+), or in the global brain inflammation state, based on the expression of the *S100a9*, *Ngp,* or in the expression of other key brain genes, such as *Plp1* and *Dcx*, when comparing the brains of both uninfected vs IAV-infected control and pDC-less mice (**Supplementary Fig.1c-g**). Altogether, these data exclude a key role of pDCs in intrinsic anti-viral immunity and in the potential neuropathy induced during pulmonary IAV infection.

**Fig. 3:**
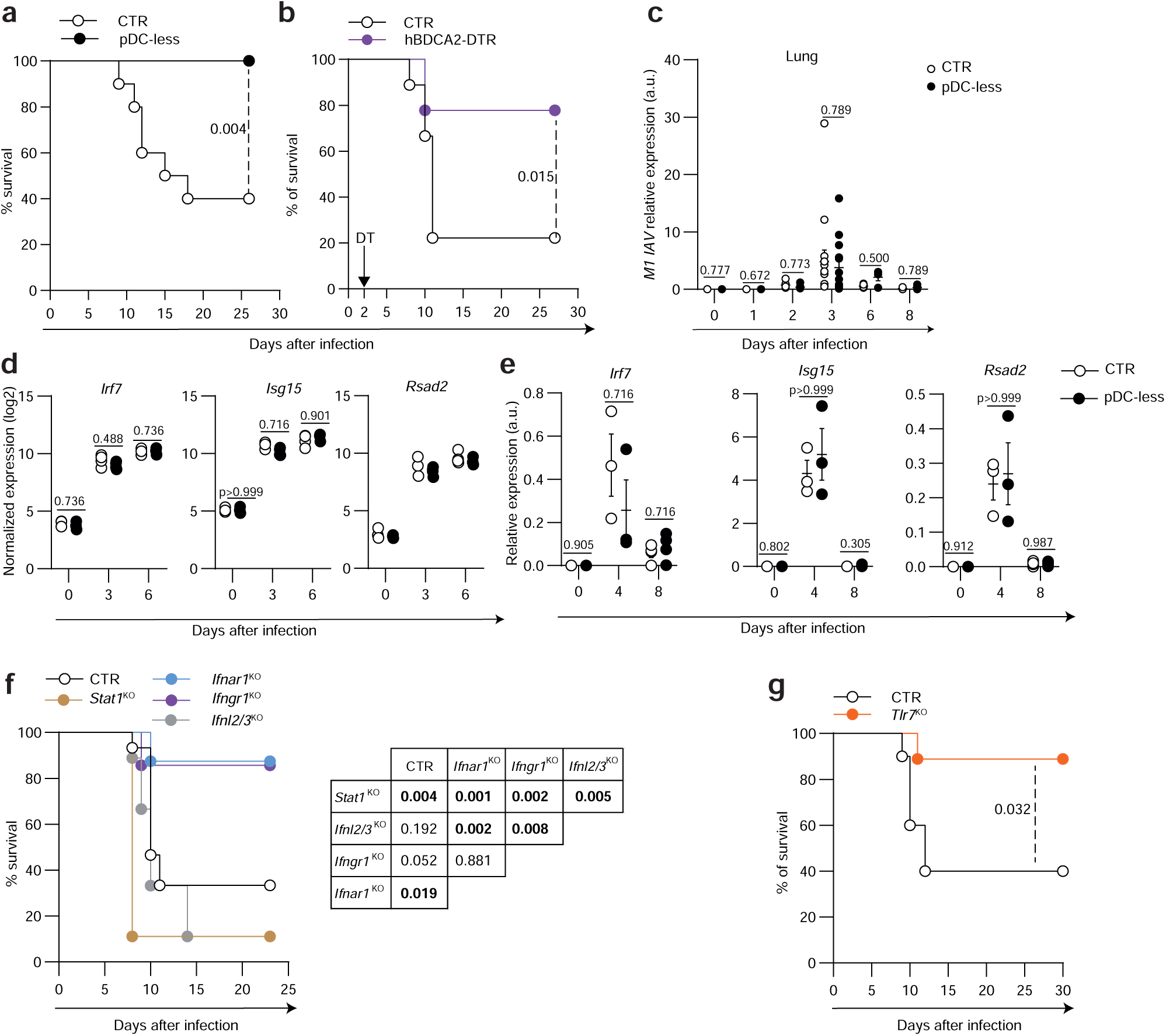
Both pDCs and the response to IFN-I are detrimental during IAV infection. Mice were infected with 145 PFU (i.n). of IAV strain H3N2A/Scotland/20/74. **a-b**, IAV-infected mice were monitored daily to assess their morbidity and survival. **b**, hBDCA2-DTR mice were inoculated with 15 ng/g of diphtheria toxin (DT) two days after IAV infection. **a**, n= 10 for both strains, **b**, n=9 for both strains. Kaplan-Meier survival plot shows the percentages of survival of indicated mouse strains. **c-e**, Expression levels of IAV M1 gene (**c**) or of indicated ISGs (**e**) was analyzed by RT-qPCR in lungs isolated from uninfected or IAV-infected CTR and pDC- less mice at indicated days after infection. Data were normalized to expression level of *Actin* gene. The data (mean + s.e.m.) shown are from two independent experiments (n=4 for each condition at each time points). **c**, n= 6 at 0, 3 at day 1, 5 at day 2, 14 at day 6 and 3 at day 8 after infection, **e**, n=5 at 0, 3 at 4 and 4 at 8 days after infection. **d**, Normalized expression (log2) of indicated genes based on Bulk RNASeq samples obtained by whole lung samples isolated from CTR and pDC-less mice at day 0 (n=3/strain), 3 and 6 (n=4/strain) after IAV infection. **f-g**, IAV-infected mice were monitored daily to assess their morbidity and survival. **f**, n= 15 CTR, 8 *Ifnar1*^KO^, 7 *Ifngr1*^KO^, 9 *Ifnl2/3*^KO^ and 9 *Stat1*^KO^ mice, **g**, n= 10 CTR, 9 *Tlr7*^KO^ mice. Kaplan-Meier survival plot shows the percentages of survival of indicated mouse strains. Log-rank (Mantel-Cox) test was used for statistical analysis in **a-b**, **f-g**.

In most murine strains, including C57BL/6, Mx1, the main virulence restriction factor against IAV, is defective, due to a genetic loss-of-function mutation. Congenic C57BL/6 mice expressing at least one functional *Mx1* allele become resistant to IAV infection, even at high viral loads^43^. This resistance was proposed to be dependent on the response to IFN-I, downstream of the Tlr7-Myd88 signaling pathway^43^. Lung viral titers were increased in IAV-infected *Mx1*^+^ mice depleted of BST2^+^ cells, leading to the hypothesis that pDCs are a source of protective IFN-I in *Mx1*^+^ mice^43^. Thus, we used our pDC-less mice to determine whether their selective pDC depletion could reveal a beneficial role of pDCs in C57BL/6 mice expressing a functional Mx1. To this aim, we bred our *Pacsin1*^LSL-DTA^ mice with B6.A2G-*Mx1*^+/+^ animals, obtaining *Pacsin1*^LSL-DTA^;*Mx1*^+/+^ mice, which we then crossed with *Siglech*^iCre^ mice to generate *Mx1*^+^ pDC-less mice. B6.A2G-*Mx1*^+/+^ mice were also bred with *Ifnar1*^KO^ mice, generating *Mx1*^+^;*Ifnar1*^KO^ animals. Surprisingly, B6.A2G-*Mx1*^+^, *Mx1*^+^ pDC-less and *Mx1*^+^;*Ifnar1*^KO^ mice, all survived the infection with high viral loads of IAV (10^^6^ pfu) and did not show any significant difference in weight loss (**Extended Data Fig. 3c**), despite a trend towards increased viral titers in *Mx1*^+^;*Ifnar1*^KO^ mice at day 2 p.i. (**Extended Data Fig. 3d**). Altogether, these data exclude a critically protective role of pDCs and of IFN-I-responsiveness in *Mx1*^+^ C57BL/6 mice infected with the Scotland IAV strain. We then investigated the role of type I, II and III IFNs in the resistance to IAV infection of mice on a classical C57BL/6 genetic background. Life-threatening lung immunopathology induced by pathogenic IAV strains was abrogated in *Ifnar1*^KO^ animals ^44, 45^, supporting a deleterious role of the response to IFN-I. When infected with the LD50 of H3N2 A/Scotland/20/74 strain, 90 % of *Ifnar1*^KO^ and of *Ifngr1*^KO^ mice survived the infection, whereas most of the *Stat1*^KO^ and *Ifnl2/3* ^KO^ mice succumbed (**Fig. 3f** and **Extended Data Fig. 3e**), highlighting the detrimental role of both IFN-I and IFN-γ and the protective functions of IFN-III. In a previous study, the administration of synthetic Tlr7 antagonists was shown to inhibit IAV-dependent lung immunopathology in 129S7 mice infected with the X31 IAV strain, thus suggesting that deleterious IFN-Is were mainly produced via a Tlr7-dependent pathway in this model^46^. Consistent with this hypothesis, most of the *Tlr7*^KO^ C57BL/6 mice infected with IAV H3N2 A/Scotland/20/74 survived (**Fig. 3g** and **Extended Data Fig. 3f**).

In conclusion, the infection of C57BL/6 mice with the pathogenic H3N2 A/Scotland/20/74 IAV strain induces a life-threatening pneumonia promoted by pDCs and the response to IFN-I and IFN-γ, likely via Tlr7-dependent mechanisms, whereas host protection relies on Stat1-dependent responses, likely mediated by IFN-III.

### During IAV infection lung pDCs increase in numbers and are a major source of IFN-α

As both pDCs and IFN-Is were harmful during infection with the Scotland IAV strain, we wondered whether pDCs could be a detrimental source of these cytokines. We determined whether and when lung pDCs produced IFN-I during IAV infection, by using *Ifnb1*^EYFP^ reporter mice^47^. pDCs, as well as tDCs and cDCs, were rare within CD45^+^ cells isolated from the lungs of uninfected mice, while their percentages increased starting from day 4 after infection (**Fig. 4a** and **Extended Fig. 4a-b**). We analyzed YFP expression, correlating with IFN-I production^37, 47^, in these different DC types. Within DCs isolated from the lungs of IAV-infected *Ifnb1*^EYFP^ mice, only pDCs contained YFP^+^ cells (**Extended Fig.4a**) that reached the plateau between day 4 and 6 after infection (**Fig. 4b**), confirming pDC contribution to IFN-I production in the lungs of IAV-infected mice. We then evaluated the impact of pDC loss on the IFN-Is detectable in the lungs, at the transcriptional and protein levels. First, we analyzed the expression of genes encoding IFN-I/IIIs, in bulk RNA-seq samples derived from the lungs isolated from control vs pDC-less mice at days 0, 3 and 6 after IAV infection (**Extended Data Fig.4c**). The expression of all the genes encoding IFN-Is and IFN-IIIs was detectable exclusively in infected mice. *Ifnb1*, *Ifna4*, *Ifnl2* and *Ifnl3* were induced to comparable levels upon infection in both mouse strains, whereas we observed a tendency to a lower expression of *Ifna1, Ifna2, Ifna6* and *Ifna5* in pDC-less mice (**Extended Data Fig.4c**). We then analyzed IFN-α and IFN-β protein contents in bronchioalveolar lavages (BAL) (**Fig. 4c**). Consistent with our transcriptional data, both IFN-I subtypes were detectable in the BAL starting day 3 after infection. However, only IFN-α levels were significantly reduced in the absence of pDCs, while IFN-β levels were comparable (**Fig. 4c**). Taken together, these results confirm that pDCs are a main source of IFN-α in infected lungs, although other cells produce other IFN-I subtypes, as well as IFN-III. Breeding *Ifnb1*^EYFP^ and pDC-Tom reporter mice generated SCRIPT animals that enable discriminating IFN+ (Tom+ YFP+) pDCs from their IFN-(Tom+ YFP-) counterparts in tissues^32^. We quantified the distribution of total vs IFN+ pDCs in the lungs of IAV-infected SCRIPT mice. Heterogeneous CD45 staining allowed to distinguish lowly inflamed (CD45^low^) vs highly inflamed (CD45^high^) lung areas, the latter ones resembling inflammatory foci (**Fig. 4d-e, j** and **Extended Data Fig. 4d-e**). Highly inflamed areas contained also the highest numbers of infected cells expressing viral hemagglutinin (HA^+^) (**Fig. 4d-e, k** and **Extended Data Fig. 4f**). Both CD45^+^ and HA^+^ cell numbers increased during infection. pDC frequency significantly increased in highly inflamed areas, starting from day 4 after infection (**Fig. 4f-g, l** and **Extended Data Fig. 4g**), consistent with our flow cytometry data (**Fig. 4a** and **Extended Data Fig. 5a-b**). IFN^+^ pDCs (YFP^+^ Tom^+^) were detected only in highly inflamed areas and peaked at day 4 after infection (**Fig. 4h-i, m** and **Extended Data Fig. 4h**). Thus, during IAV respiratory infection, lung pDCs produce IFN-Is locally, preferentially in highly inflamed and infected areas.

**Fig. 4:**
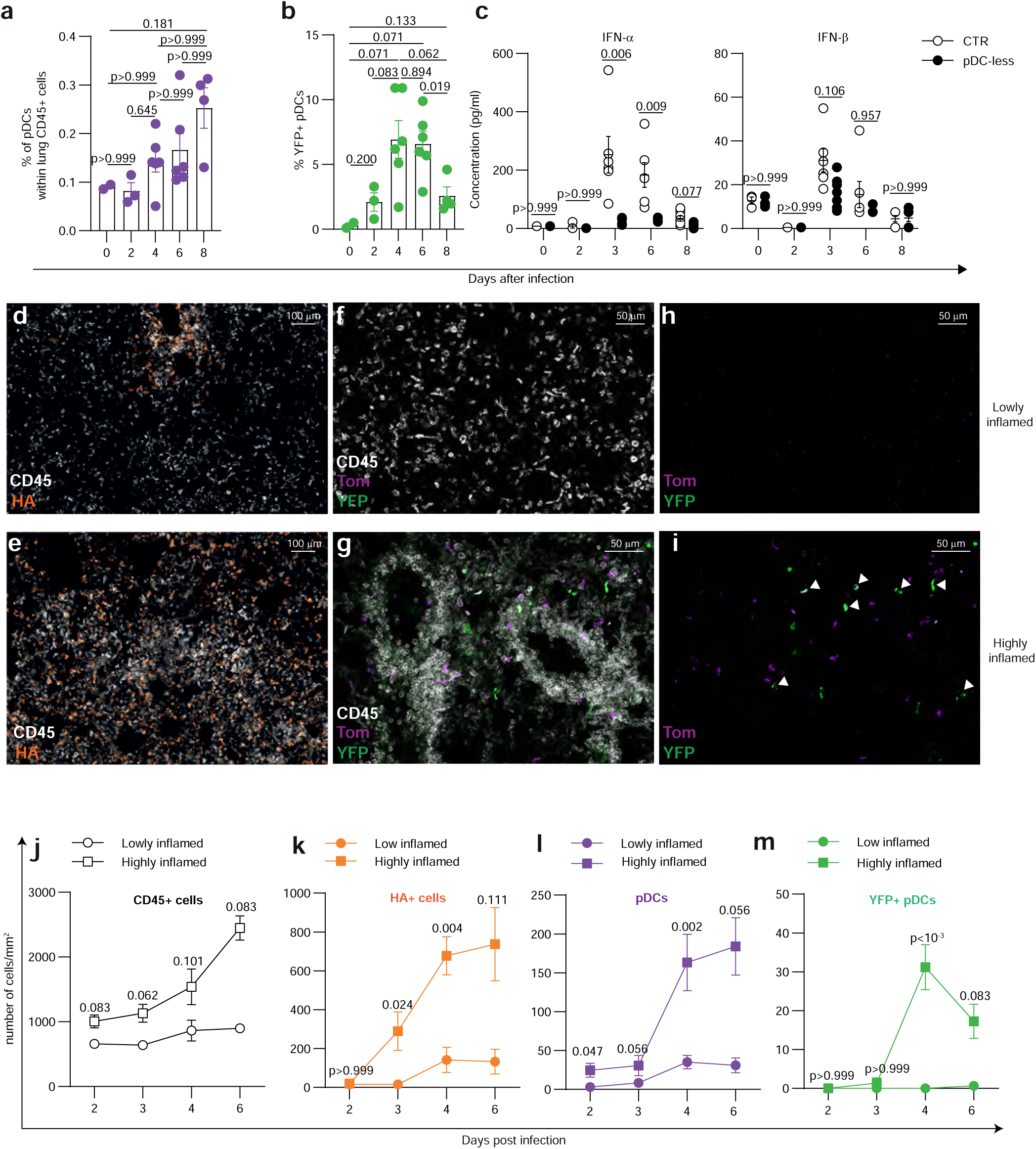
During IAV infection, pDCs increase in infected lungs and produce IFN-I. a-b, *Ifnb1*^EYFP^ mice were infected (i.n.) with 160 PFU of IAV strain H3N2A/Scotland/20/74 and analyzed at different time points after infection. **a**, The percentages of pDCs within CD45^+^ cells (mean +/- s.e.m.) isolated from the lungs of infected *Ifnb1*^EYFP^ mice were determined at the indicated time points. The percentages of YFP^+^ pDCs (**b**) at the indicated days after infection have been represented. N=2 for day 0, 3 for day 2, 6 for days 4 and 6, 4 for day 8 after infection. In **a-b**, a two-tailed non-parametric Mann-Whitney test was used for the statistical analysis. **c,** Cytometric bead arrays was performed to quantify IFNα and IFNβ concentrations in the BAL isolated from CTR and pDC-less mice uninfected (0) or at indicated days after infection. Each dot represents an individual animal. The data shown (mean +/- s.e.m.) are pooled from two independent experiments (n=3 CTR and 4 pDC-less at day 0, 3 CTR and 2 pDC-less at day 2, 7 at day 3, 6 at day 6 and 7 at day 8 for both strains). An unpaired and nonparametric multiple t-test (Mann-Whitney) with Holm-Sidak method for correction was used for the statistical analysis. **d-i**, Representative images of histological sections isolated from the lungs of SCRIPT mice at day 4 after IAV infection. 30µm-thick lung sections were stained with anti-CD45 (white), anti-Hemagglutinin (HA) of IAV (orange), anti-Tomato (Tom, purple), anti-YFP (green). On each image acquired, the MFI of CD45 in highly inflamed areas was at least twice of that of lowly inflamed areas. The numbers of CD45^+^ cells (**j**), HA (IAV)^+^ cells (**k**), pDCs (**l**) and YFP^+^ pDCs (**m**) per mm² were quantified in highly and lowly inflamed areas in IAV-infected lungs. The data shown (mean +/- s.e.m.) are from n=5 at day 2, 5 at day 3, 10 at day 4 and 4 at day 6, with two sections of the lung analyzed per mouse. An unpaired and nonparametric multiple t-test (Mann-Whitney) with Holm-Sidak correction method was used for the statistical analysis.

### IFN production by pDCs is mainly responsible for their deleterious role during IAV infection

During IAV infection, pDCs are a main source of IFN-α, but not of IFN-β, in the lung (**Fig. 4** and **Extended Fig. 5**). Studying the specific role of the IFN-I produced by pDCs in IAV-induced pneumonia would require the inactivation of genes involved in the TLR7/Myd88/IRF7 signaling cascade uniquely in pDCs. However, none of the existing conditional Cre mice, including our Siglech^iCre^ strain^32, 48^, allows the selective targeting of pDCs. To overcome this roadblock, we developed a novel chimeric mouse model, the pDC Shield Bone Marrow Chimeras (pDC SBMC) mice (**Fig. 5a**). These mice were generated by lethally irradiating the hind legs of pDC-less mice, while the rest of their body was protected by a lead shield. This promoted partial engraftment of donor BM cells preferentially reconstituting the immune cell lineages lacking in the recipient mice, in our case pDCs. Indeed, four weeks after reconstitution, pDC SBMC mice were replenished with pDCs derived from donor BM, whereas the other immune lineages were still mostly composed by recipient cells, as validated by using CD45.2 donor BM cells in CD45.1^+^ pDC-less mice that were used as SBMC recipient (**Fig. 5b**). pDC SBMC thus allow to study the impact of the loss of a candidate gene selectively in pDCs. As cytokine production by pDCs is Myd88-dependent^48^, we evaluated the resistance to IAV infection of pDC SBMC reconstituted with *Myd88*^KO^ pDC. We generated CTR, pDC-less and *Myd88*^KO^ pDC SBMC by reconstituting shield-irradiated CD45.1^+^ pDC-less mice with CD45.2^+^ C57BL/6, pDC-less or *Myd88*^KO^ BM cells, respectively. When infected with IAV, half of CTR pDC SBMC succumbed, whereas all pDC-less and most of the *Myd88*^KO^ pDC SBMC survived (**Fig. 5c** and **Extended Fig. 5a**). *Tlr7*^KO^ pDC SBMC were also resistant to IAV infection, when compared to infected CTR pDC SBMC (**Fig. 5d** and **Extended Fig. 5b**). pDC-intrinsic lack of TLR7/Myd88 completely abrogates TLR7-dependent cytokine and chemokine production, while pDC-intrinsic deficiency in IRF7 abolishes exclusively IFN production ^48^. We thus generated *Irf7*^KO^, as well as CTR, pDC-less pDC SBMC and infected them with IAV. Most of the *Irf7*^KO^ and pDC-less pDC SBMC survived, whereas 60% of the CTR pDC SBMC died (**Fig. 5e** and **Extended Fig. 5c**). Consistently, *Irf7*^KO^ and pDC-less pDC SBMC displayed overall reduced morbidity and clinical scores based on their posture, mobility, fur appearance and respiration, when compared to CTR pDC SBMC (**Extended Fig. 5d**). However, histological analysis of the lungs of the different SBMC did not show any significant difference in lesion scoring (**Extended Fig. 5e**). *M1* IAV expression was comparable in CTR, *Irf7*^KO^ and pDC-less pDC SBMC (**Fig. 5f**), showing similar viral loads. *Ifna1*, *Ifna2* and *Ifna5* expression was abrogated in IAV-infected *Irf7*^KO^ and pDC-less pDC SBMC, as compared to CTR, while *Ifna4* and *Ifnb1* were similarly induced in the three types of SBMC (**Fig. 5f**). Moreover, at the protein level, the levels of IFN-α, but not of IFN-β, were significantly decreased in the BAL of IAV-infected *Irf7*^KO^ and pDC-less pDC SBMC (**Fig. 5g**). These results are thus reminiscent of the observations in IAV-infected pDC-less mice (**Fig.4** and **Extended Fig.4**).

**Fig. 5:**
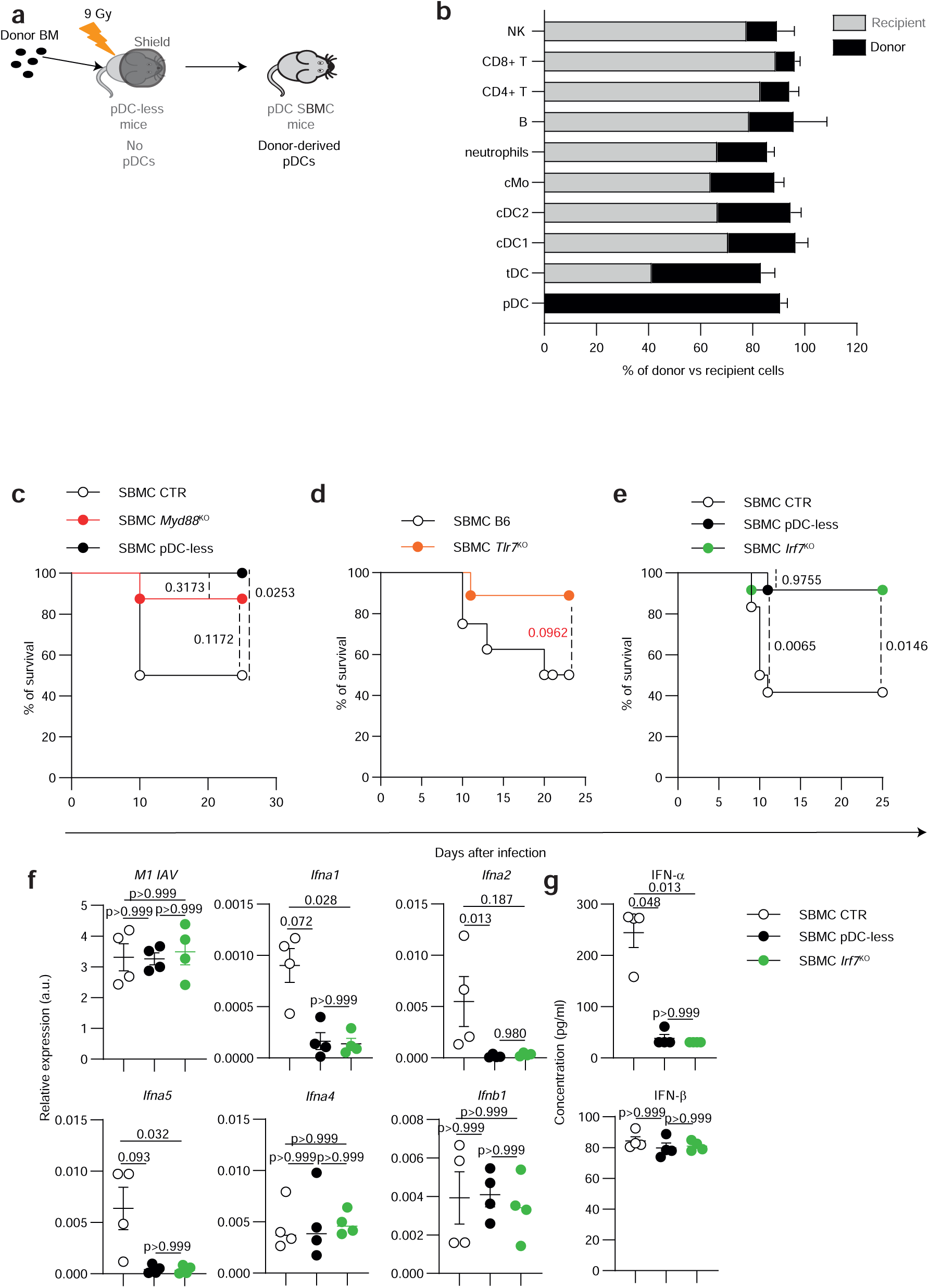
IFN-I produced by pDCs during IAV infection are deleterious. a, Scheme illustrating the generation of pDC Shield BM chimeric (SBMC) mice. Only hind limbs of pDC-less mice are irradiated, while the rest of the body is protected by a lead shield (gray oval). Upon local irradiation-induced distress, donor BM cells preferentially reconstitute immune cell types absent in the recipient, that is pDCs. **b**, Recipient CD45.1+ (gray) pDC-less mice are used as recipient of pDC SBMC reconstituted with donor CD45.2+ (black) BM cells. The percentage of donor (black) vs recipient (gray) cells was analyzed in indicated lineages 4 weeks after reconstitution (n= 4). **c-e,** pDC SBMC mice reconstituted with CTR, pDC-less or *Myd88*^KO^ BM (**c**), with CTR or *Tlr7*^KO^ BM (**d**) or with CTR, pDC-less or *Irf7*^KO^ were infected (i.n). with 145 PFU of IAV strain H3N2A/Scotland/20/74 IAV-infected mice and daily monitored to assess their morbidity and survival. Kaplan-Meier survival plots show the percentages of survival of indicated genotypes. **c-d,** Pool of two independent experiments (n=8 for each strain); **e,** Pool of three independent experiments (n=12 for each strain). Log-rank (Mantel-Cox) test was used for statistical analysis in **c-e**. **f**, Expression of indicated genes was analyzed by RT-qPCR in lungs isolated from indicated IAV-infected SBMC 3 days after infection. **g**, Cytometric bead arrays was performed to quantify IFNα and IFNβ concentrations in the BAL isolated from the same SBMC used in **f**. Each dot represents an individual animal. n=5 in **f-g**.

In conclusion, selective inactivation of Tlr7/Myd88/Irf7 signaling in pDCs restored resistance to IAV infection in most of the pDC SBMCs and ameliorated their clinical score, thus supporting the detrimental contribution of the IFNs produced by lung pDCs during IAV-induced lung immunopathology.

### During IAV infection, lung inflammation and cytokine contents are reduced in the absence of pDCs

During IAV infection, the recruitment of immune cells to the infected lungs and their activation are essential to establish protective anti-viral immunity. However, unbridled and sustained inflammatory responses can lead to severe pneumonia and acute respiratory distress syndrome (ARDS). Therefore, we performed anatomopathological analysis of lung sections isolated from IAV-infected control and pDC-less mice, to evaluate the impact of pDC loss on lung inflammation and tissue integrity. We harvested our samples 8 days after IAV infection, when survival and morbidity start to differ between these two mouse strains (**Fig. 3a**). Most of the lungs isolated from infected control mice showed a marked broncho-interstitial pneumonia (**Fig. 6a**), eventually associated with bronchiolar filling by degenerated epithelial cells and with intense alveolar inflammation (**Fig. 6b**). In contrast, lungs isolated from infected pDC-less mice displayed milder pneumonia and inflammation (**Fig. 6c-d**). Consistently, semi-quantitative score of lung sections, taking in account inflammation in interstitial, alveolar, bronchial and vascular areas (**Suppl Table 1**), was significantly higher in control than in pDC-less mice (**Fig. 6e**). In a previous study, in IAV-infected 129S7 mice infected with the IAV X31 strain, the depletion of BST2^+^ cells correlated with reduced recruitment of inflammatory leukocytes in infected lungs^44^. In our study, the absolute numbers of whole lung cells and, especially, of CD45^+^ cells, increased during infection and peaked at 6 days p.i., following a similar trend in infected control and pDC-less mice (**Fig. 6f** and **Extended Fig. 7a-c**). The percentages of most myeloid and lymphoid lineages isolated from infected control and pDC-less mice were comparable, except for pDCs, as expected (**Fig. 6g** and **Extended Data Fig. 7a-d**). However, both neutrophils and eosinophils were significantly increased at days 6-8 after IAV infection in the absence of pDCs (**Fig. 6g**). Neutrophils were proposed to exert detrimental vs protective roles upon IAV infection^46, 49^. We thus investigated whether neutrophils were required for enhanced resistance of pDC-less mice to IAV infection, by injecting animals with isotype control (IC) or neutrophil-depleting antibodies (1A8) at days 5 and 6 after infection. Neutrophil depletion severely affected the survival of both control and pDC-less infected mice (**Figure 6h**), showing that, in our infection model, neutrophils were protective, but via pDC-independent mechanisms. Indeed, selective pDC depletion did not inhibit the recruitment of immune cells into infected lungs; rather it was associated with neutrophilia and eosinophilia.

**Fig. 6:**
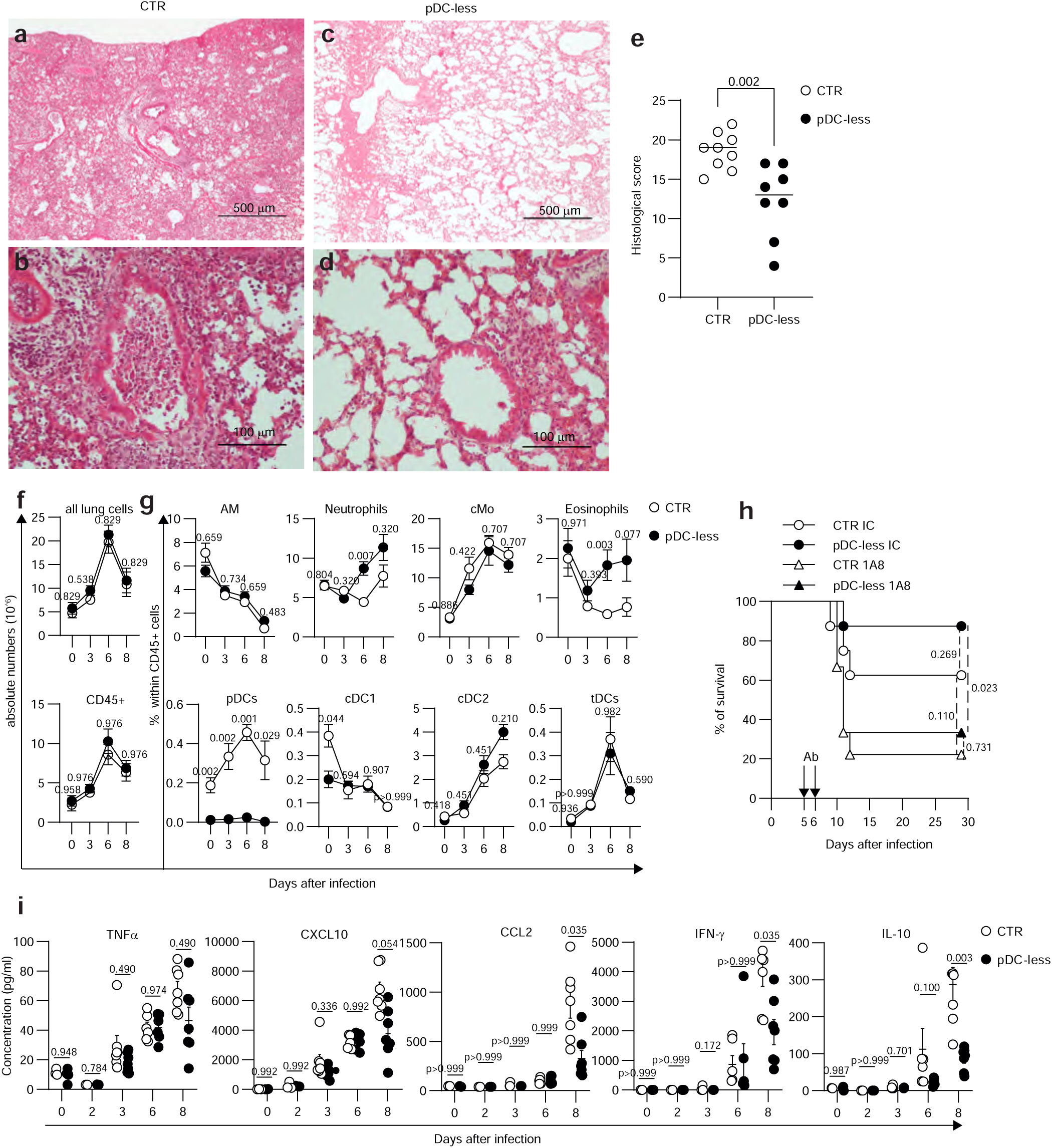
During IAV infection, lung inflammation and cytokine production are reduced in the absence of pDCs. a-d, Representative images of H&E-stained sections of lungs of control (**a-b**) and pDC-less (**c-d**) mice isolated at day 8 after infection. (**a-b**) marked broncho-interstitial pneumonia with terminal bronchiole filled with degenerated neutrophils; (**c-d**) mild broncho- interstitial pneumonia. Magnification is indicated on each image. **e,** Lung sections of all mice were semi-quantitatively assessed using a cumulative score as normal (0), mild (1), moderate (2), marked (3) or severe (4) for inflammation in interstitium, intra-alveolar, peribronchial, intra- bronchial and perivascular areas, and as normal (0), focal (1), multifocal (2), extensive (3) for hemorragic necrosis, emphysema and pleura. The data (mean +/- s.e.m.) shown are pooled from two independent experiments, n=8 CTR, 9 pDC-less mice. A two-tailed nonparametric Mann-Whitney test was used for the statistical analysis. **f,** Absolute numbers of all lung cells and of CD45^+^ cells isolated from the lungs of control and pDC-less mice at the indicated days after infection. Data are represented as mean +/- s.e.m. of the following numbers of individuals pooled from two independent experiments for each time point (n= 5 at day 0, 7 at day 3, 8 at day 6 and at day 8 for each mouse strain). **g,** Percentages of myeloid and lymphoid immune cells isolated from the lungs of control and pDC-less mice isolated at the indicated day after infection. n= 5 at day 0, 7 at day 3, 8 at day 6 and 4 at day 8, except n= 8 at day 8 for neutrophils, eosinophils, alveolar macrophages (AM) and monocytes (cMo). For **f** and **g**, an unpaired and nonparametric multiple t-test (Mann-Whitney) with Holm-Sidak correction method was used for the statistical analysis. **h,** Control and pDC-less mice were infected with 145 PFU of IAV strain and treated with anti-Ly6G (1A8) antibody (triangle) or isotype control (IC,circle) at days 5 and 6 after infection. Survival and morbidity were daily monitored. Data are pooled from two independent experiments. Kaplan-Meier survival plot shows percentage survival of indicated conditions. For each mouse strain, n= 8 for mice treated with isotype control (IC) and n=9 for mice treated with 1A8 antibodies. Log-rank (Mantel-Cox) test was used for statistical analysis. **i,** Cytometric bead arrays was performed to quantify the concentrations of indicated cytokines in the BAL isolated from control and pDC-less mice at indicated days after IAV infection. Each dot represents an individual. Data shown (mean + s.e.m.) are pooled from two experiments. n= 3 CTR and 4 pDC-less mice at day 0, n= 3 CTR and 2 pDC-less mice at day 2, n= 7 at day 3, 6 at day 6 and 7 at day 8 for both mouse strain. An unpaired and nonparametric multiple t-test (Mann-Whitney) with Holm-Sidak method for correction was used for the statistical analysis.

**Fig. 7:**
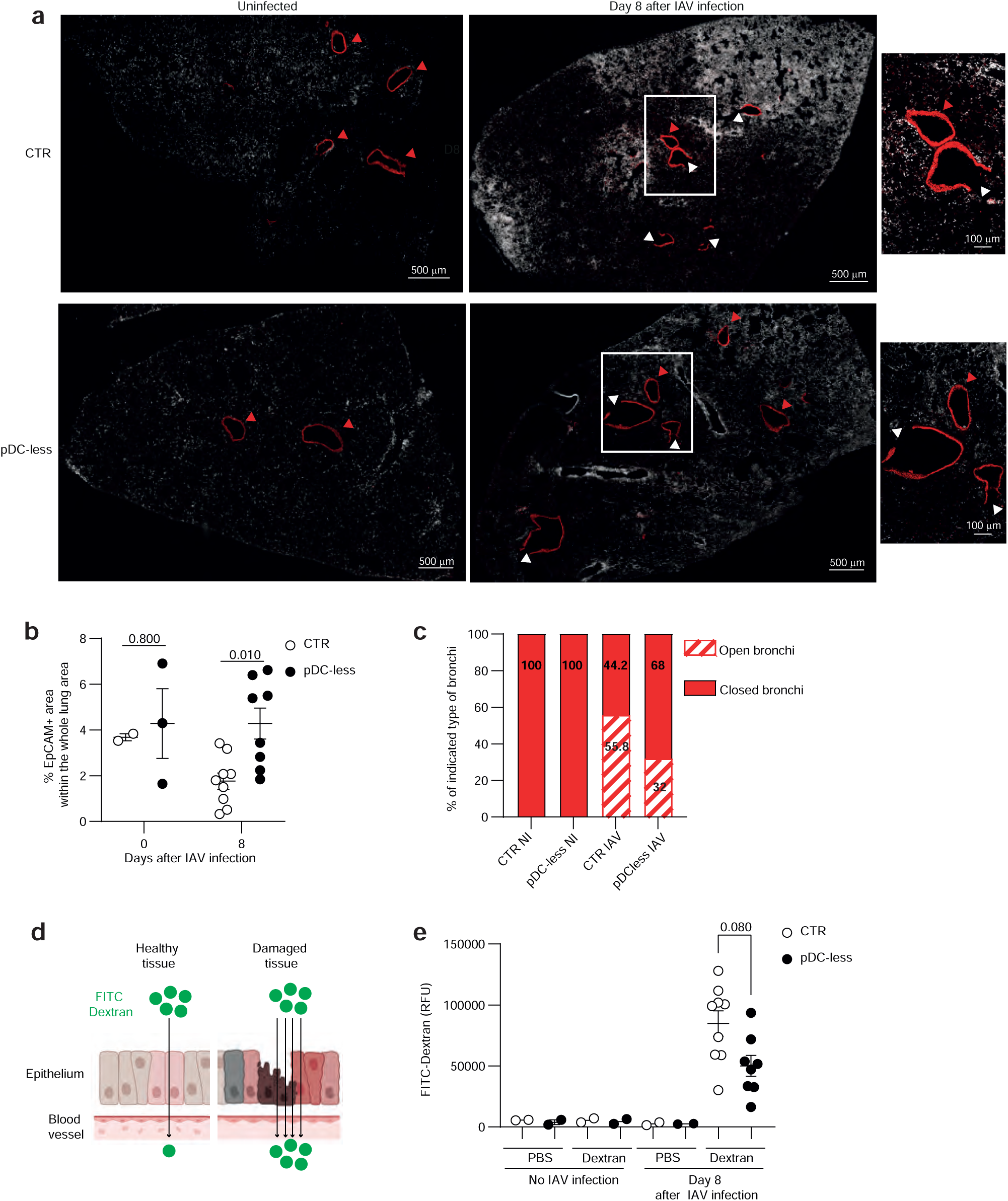
During IAV infection lung integrity and permeability are less affected in the absence of pDCs. **a**, Representative images of histological sections of lungs isolated from control and pDC-less mice uninfected (left) or at day 8 after IAV infection (right). 30µm-thick sections were stained with anti-CD45 (white) and anti-EpCAM (red) antibodies. EpCAM+ bronchi were classified as closed (red arrow) when EpCAM staining was uninterrupted and as opened (white arrow) when EpCAM was discontinuous. **b-c**, The percentages of the EpCAM+ area over the entire surface of each lobe (**b**) and the percentages of closed vs opened bronchi (**c**) were quantified in lung sections isolated from uninfected vs IAV-infected control vs pDC- less mice. The data (mean +/- s.e.m.) shown in **b-c** are from n=1 at day 0 and n=4 at day 8 after infection for both mouse strains. We analyzed 2 lobe sections at day 0 and 9 lobe sections at day 8 for control mice, 3 lobe sections at day 0 and 8 lobe sections at day 8 for pDC-less. An unpaired and nonparametric multiple t-test (Mann-Whitney) with Holm-Sidak correction method was used for the statistical analysis. **d-e,** 8 days after IAV infection control and pDC- less mice were i.t. treated with fluorescein isothiocyanate (FITC-Dextran) (10µg per mouse). Barrier permeability was measured as relative fluorescent units (RFU) of FITC-dextran leaked in the plasma 1 hour after i.t. FITC-dextran administration. Each dot represents an individual. Data shown (mean + s.e.m.) are pooled from two experiments (n = 2 for each strain and condition for uninfected mice, as well as for PBS-treated infected mice, n= 9 for FITC-treated infected CTR and 8 for pDC-less mice). An unpaired and nonparametric multiple t-test (Mann-Whitney) with Holm-Sidak method for correction was used for the statistical analysis.

IFN-Is produced by pDCs activate both innate and adaptive immune cells^1^, by inducing the expression of activation molecules, such as CD86 and CD69. The expression of both these markers was induced during infection on lung myeloid and lymphoid cells, but it was only transiently affected in the absence of pDCs (**Extended Data Fig. 8a-b**). In a previous study, the depletion of BST2^+^ cells was also shown to reduce the amounts of proinflammatory cytokines detected in bronchioalveolar lavages (BAL) of 129S7 mice infected with the IAV X31 strain, and, consequently, lung immunopathology^44^. The analysis of our bulk RNA-seq data showed that mRNAs encoding most of the cytokines and chemokines analyzed were similarly upregulated in infected control and pDC-less mice at days 3 and 6 after infection (**Extended Data Fig. 8c**). BAL isolated from both strains of mice at days 2, 3 and 6 after infection did not differ in their content for proinflammatory cytokines, including TNF, CXCL10, CCL2, IFN-γ, IL-6 and CCL5, and anti-inflammatory cytokines, such as IL-10 (**Fig. 6i** and **Extended Data Fig. 8d**). However, at day 8 after infection, the amounts of CXCL10, CCL2, IFN-γ and IL-10 detectable in the BAL were significantly lower in the absence of pDCs (**Fig. 6i** and **Extended Data Fig. 8d**). In contrast, TNF, IL-6 and CCL5 levels were not affected, suggesting pDC-independent sources. Hence, during late phases of IAV infection, enhanced resistance of pDC-less mice correlated with reduced bronchioalveolar amounts of some of the cytokines known to promote ARDS.

### The alterations of lung integrity and permeability induced upon IAV infection are less severe in the absence of pDCs

Unbridled cytokine-dependent inflammation induced by IAV infection can promote severe lung lesions, fluid leakage and defaults in respiratory capacity, leading to ARDS. Reduced cytokine levels in the BAL of IAV-infected pDC-less mice could indicate reduced tissue damage and leakage in the bronchial tree in the absence of pDCs, consistent with milder histopathological score in the lungs of infected pDC-less mice (**Fig. 6e**). Bronchi were clearly identified in the lung sections of uninfected control and pDC-less mice by staining with antibodies recognizing Epithelial Cell Adhesion Molecule (EpCAM), a marker of bronchiolar cells (**Fig. 7a**). However, whereas in uninfected lungs EpCAM staining was continuous and homogeneous in both strains (**Fig. 7a**, red arrows), it was often disrupted in the lungs of infected mice (**Fig. 7a**, white arrows). On the whole lung sections, the percentages of EpCAM^+^ areas were significantly higher in infected pDC-less mice (**Fig. 7b**), whereas the percentages of bronchi with disrupted (open) EpCAM staining were higher in infected control mice (**Fig. 7c**). These data suggested an increased breaching of the bronchial tree, and, potentially, a higher fluid leakage, in the presence of pDCs. Pulmonary permeability can be analyzed by measuring the leakage into the plasma of fluorescent proteins instilled in the airways (**Fig. 7d**)^50^. We thus instilled 10KDa FITC-Dextran intratracheally in control and pDC-less mice, uninfected or 8 days after IAV infection. While the fluorescence was absent in the plasma of uninfected mice, as well as of PBS-instilled IAV-infected mice, it was clearly detectable in the plasma of IAV-infected animals and significantly lower in pDC-less than in control mice (**Fig. 7e**). Thus, pDCs contribute to promote lung lesions and fluid leakage during IAV infection.

### During mouse SARS-CoV2 infection, pDCs are not required for host survival and can exert detrimental functions

Patients harboring loss-of-functions genetic mutations in *MYD88*, *IRAK4, IRF7* or *TLR7* were highly susceptible to the infection by several respiratory viruses, including Influenza and SARS-CoV2^13, 14, 15, 16, 17^. As blood pDCs of these patients were defective in vitro in TLR7-dependent IFN production, patient susceptibility was attributed to impaired pDC-dependent viral control^14, 17^. However, other studies proposed rather a detrimental role of pDCs during SARS-CoV2 infection, through an IFN-I-dependent activation of macrophages fueling a cytokine storm^51, 52, 53, 54^. We thus used our pDC-less mice to investigate the specific role of pDCs in SARS-CoV2 infection. C57BL/6 mice typically do not develop detectable infection or pathology when exposed to most SARS-CoV2 strains, largely because these viral strains exhibit low affinity binding to mouse angiotensin-converting enzyme 2 (ACE2), the primary receptor for viral entry in the cells. However, ectopic expression of human ACE2 under the control of the cytokeratin 18 (K18) promoter in transgenic K18:hACE C57BL/6 mice render them highly susceptible to SARS-CoV infections^55^. Hence, we first generated K18:hACE x *Pacsin1*^LSL-DTA^ mice, then we bred them with *Siglech*^iCre^ mice. Offspring triple heterozygous mice, hereafter called K18:hACE pDC-less mice, as well as control K18:hACE mice, were infected with the SARS-CoV2 Wuhan/D614 strain. Infected K18:hACE males survived less than females, consistent with the higher morbidity and mortality observed in male SARS-CoV2-infected patients^56^ (**Fig. 8a** and **Extended Data Fig. 8a**). We did not detect any significant difference in survival between infected female K18:hACE mice and K18:hACE pDC-less mice. In contrast, infected males K18:hACE pDC-less mice were significantly more resistant than gender-matched infected K18:hACE mice (**Fig. 8a** and **Extended Data Fig. 8a**). Thus, pDCs are not required for K18:hACE mouse survival to SARS-Cov2 infection and appear to exert detrimental functions in infected males.

**Fig. 8:**
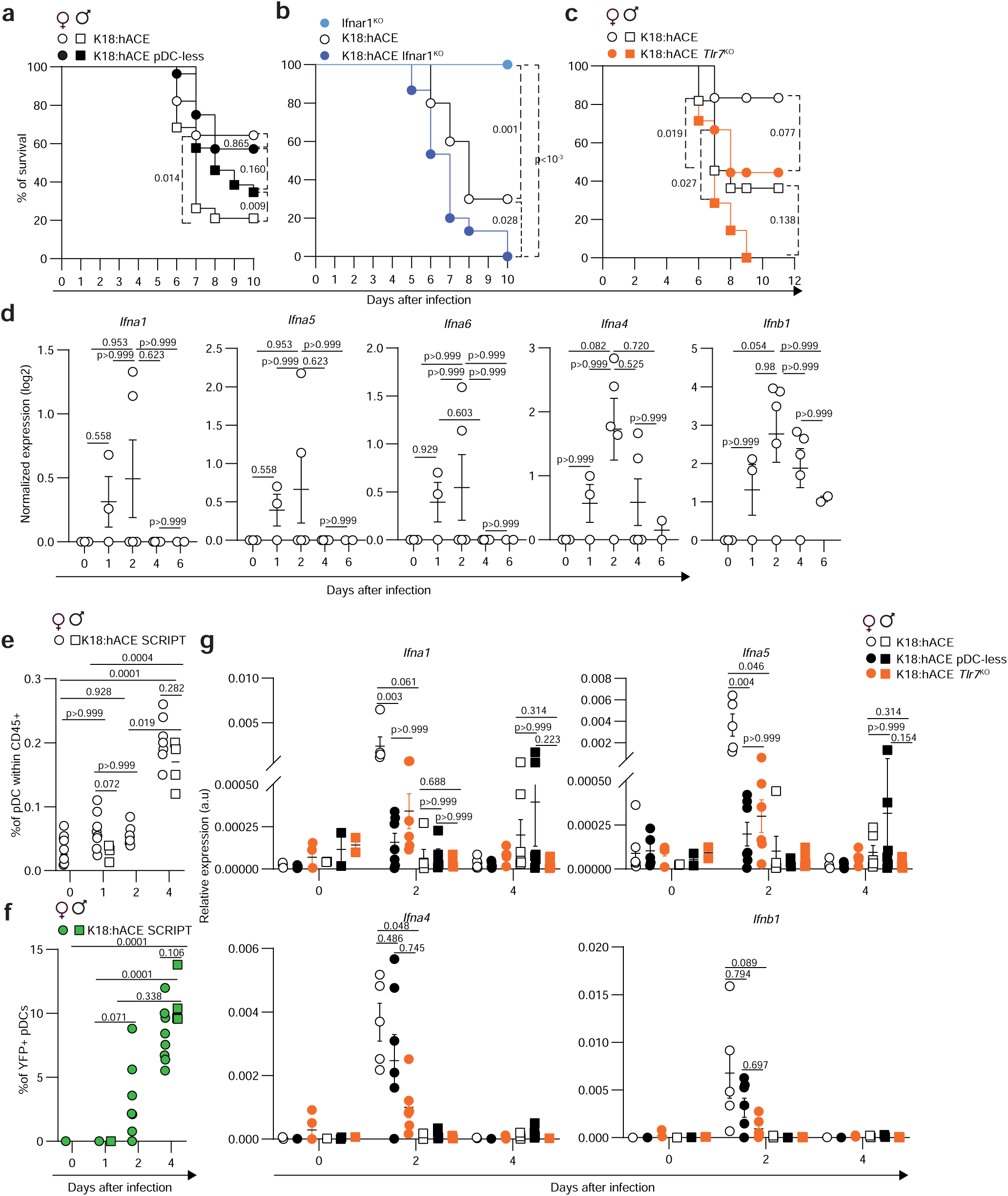
**pDCs are not required for the resistance to SARS-CoV2 infection and can exert detrimental functions**. **a-c,** K18:hACE transgenic mice bred or not with indicated mutant mice were infected with 200 PFU (i.n). of SARS-CoV2 strain. Infected mice were daily monitored to assess their morbidity and survival. Kaplan-Meier survival plot shows the percentages of survival of indicated mouse strains. **a**, n= 28 for female K18:hACE and K18:hACE pDC-less; 19 for male K18:hACE, 26 for male K18:hACE pDC-less, **b**, n=10 for K18:hACE and plain *Ifnar1*^KO^ 15 for K18:hACE *Ifnar1*^KO^, **c**, n= 12 for female K18:hACE, 9 for female K18:hACE *Tlr7*^KO^; 11 for male K18:hACE, 7 for male K18:hACE *Tlr7*^KO^. Log-rank (Mantel-Cox) test was used for statistical analysis. **d**, Normalized expression (log2) of indicated genes based on Bulk RNASeq samples obtained by whole lung samples isolated from transgenic K18:hACE mice infected with 1.1 10^5^ PFU of SARS-CoV2 at indicated days after infection. Data (mean +/-s.e.m.) shown are from n= 3 at days 0 and 1, 5 at day 2 and 4, 2 at day 6 after infection. **e-f**, Percentages of pDCs (**e**) and YFP+ pDC (**f**, IFN-producing pDCs) within CD45^+^ lung cells isolated from K-18:hACE SCRIPT infected with 1.1 10^5^ PFU of SARS-CoV2 at indicated days. n= 10 females (F) at 0, 8 F and 4 males (M) at 1, 7 F at 2 and 9 F and 4 M at 4 days p.i. **g**, Expression levels of indicated genes were analyzed by RT-qPCR in lungs isolated from uninfected or SARS-CoV2-infected control (CTR), pDC-less and *Tlr7*^KO^ K18:hACE mice at indicated days after infection. Data were normalized to expression level of *Actin* gene. The data (mean + s.e.m.) shown are from two independent experiments (day 0= 7F and 2M CTR, 5F and 2M pDC-less, 4F and 2M *Tlr7*^KO^, day 2= 5F and 5M CTR, 7F and 5M pDC-less, 6F and 5M *Tlr7*^KO^, day 4= 6F and 6 M CTR for each strain). A nonparametric one-way ANOVA (Kruskal-Wallis test followed by Dunn’s post-hoc test) was used for the statistical analysis in **d-g**.

The response to IFN-I is essential for human survival to many viral infections, including to SARS-CoV2^15^. However, IFN-I-responsiveness can be deleterious in IAV infected-mice^44, 45, 46^ (**Fig. 3** and **Fig. 5**). When infected with SARS-CoV2, female K18:hACE *Ifnar1*^KO^ mice were more susceptible than female K18:hACE (**Fig. 8b** and **Extended Data Fig. 8b**), thus supporting a main protective role of IFN-I in this infection model. In contrast, plain *Ifnar1*^KO^ mice were resistant, consistent with their lack of the hACE receptor. Viral loads were not different between control, pDC-less and *Ifnar1*^KO^, K18:hACE mice (**Extended Data Fig. 8c**), suggesting that the differences in survival between these mouse strains were mostly due to differences in immunopathology rather than in viral control. Patients with TLR7 mutations, especially males, were found highly susceptible to SARS-CoV2 infection^16^. We thus addressed the role of TLR7 in K18:hACE mice, by breeding them with *Tlr7*^KO^ mice. TLR7 deficiency significantly increased the susceptibility of K18:hACE mice to SARS-CoV2 infection, independently of mouse gender, demonstrating a protective role of TLR7 (**Fig. 8c** and **Extended Data Fig. 8d**). Altogether, these results demonstrate that the survival of K18-hACE mice to SARS-CoV2 infection requires a TLR7-dependent IFN-I production by other cell types than pDCs.

The timing and the magnitude of IFN production during SARS-CoV2 infection varies between patients developing moderate versus severe COVID^51^. In mice infected with SARS-CoV1/2 and MERS-CoV2, IFNs were beneficial in very early phases of the infection, but detrimental later on^57, 58, 59^. In our mouse model, the expression of IFN-I and ISGs in SARS-CoV2-infected lungs was induced as soon as 1-2 days after infection, consistent with SARS-CoV2 detection in infected tissues (**Fig. 8d-e** and **Extended Fig. 9a-b**). The induction of ISGs persisted after the peak of IFN expression (day 4), and was not affected in infected K18:pDC-less mice (**Extended Fig. 9a-b**), according with their dispensable role for viral control. In contrast, ISG upregulation was affected to variable extents in infected K18:hACE *Ifnar1*^KO^ mice (**Extended Fig. 9a-b**), likely due to partial compensation by other IFNs, such as IFN-II and IFN-III. We then analyzed the kinetics of pDCs and their IFN production, by infecting K18:hACE bred with SCRIPT mice. Lung pDCs increased starting from day 2 after infection (**Fig. 8e and Extended Fig. 9c-d**). Similarly, YFP^+^ pDCs were detectable at day 2 and increased at day 4 (**Fig. 8f**). Thus, the peak of pDC-dependent IFN production occurs between day 2 and day 4 after SARS-CoV2 infection, likely delayed with respect to ISG induction and persisting during late phases of the infection, potentially contributing to IFN-dependent detrimental functions, as proposed in humans^54^.

As the protection of K18:hACE mice from SARS-CoV2 infection required IFN-I and TLR7, whereas pDC were dispensable or detrimental depending on host gender, we investigated the impact of the loss of pDCs versus of TLR7 signaling on the production of, and responses to, IFN-I. Viral titers were comparable in all strains and independently of host gender (**Extended Fig.9e**). In infected females, the expression of the *Ifna* and *Ifnb1* genes was induced at day 2 p.i., but barely detectable at day 4 p.i. (**Fig. 8g**). *Ifn*s were poorly detectable in infected males, with delayed expression at day 4 p.i. only in some individuals. Interestingly, in infected females, the expression of all *Ifn* genes analyzed was affected in a Tlr7-dependent manner, while only *Ifna1* and *Ifna5* appeared to be also pDC-dependent. This would suggest that, as already shown during IAV infection, pDC mainly contribute to the production of IFN-α, but not of IFN-β, and other cells than pDCs contribute to the Tlr7-dependent induction of IFN-Is. We then analyzed the expression of the ISGs *Isg15*, *Irf7* and *Rsad2* in uninfected vs SARS-CoV2 infected control, pDC-less and *Tlr7*^KO^ K18-hACE mice. In infected female mice, at day 2 p.i., the expression of all the ISGs tested was dependent on both Tlr7 and pDC to a seemingly lesser extent (**Extended Fig.9f**). We observed a similar trend, although not significant, for infected males at the same time point.

In conclusion, during SARS-CoV2 infection of K18:hACE mice, pDCs were dispensable in females and detrimental in males, whereas both IFN-I and Tlr7 were protective regardless of host gender.

## Discussion

In this study, we reported the generation of a new mouse model, the pDC-less mice, allowing specific and constitutive ablation of pDCs. pDC-less mice developed normally and were healthy, thus excluding a critical role for pDCs in host fitness under homeostatic conditions, at least in an environment free of specific pathogens. We did not detect any major disruption of other immune cell types in our mice, consistent with our intersectional genetic approach specifically targeting late pDC-committed precursors without affecting other lineages^32^. Only tDC^lo^ were slightly but significantly reduced in pDC-less mice, consistent with the identification of a shared pDC/tDC-committed bone marrow precursor^29^. Hence, in pDC-less mice, the specificity of pDC depletion is highly improved compared to previous methods that targeted pDCs but also affected DC precursors, tDCs and subsets of monocytes/macrophages or B cells^21^, including the administration of anti-PDCA1/BST2 antibodies, as well as the use of other mutant mice such as CD11c-Cre;Tcf4^-/fl^ (CKO) animals^25, 29^ and *Siglech*-hDTR animals^23, 24^. In addition, pDC-less mice allow evaluating the long-term impact of pDC loss, without the confounding side-effects caused by repeated DT injections, as observed in hBDCA2-DTR mice^30, 31^. Moreover, the use of pDC-less mice as recipient for the generation of shield BM chimeras allowed to inactivate candidate genes selectively in pDCs, for the first time to our knowledge. This thus represents an excellent surrogate method to overcome the current lack of pDC-specific Cre mice.

pDCs are highly conserved in vertebrates, both in their molecular make-up^60^ and in their functional specialization, namely professional production of IFNs, with rapid and massive release of all subtypes of these cytokines upon exposure to virus-type stimuli^1, 48^. This implies that pDC functions benefit vertebrates in a manner enhancing their reproductive fitness. The beneficial role of pDCs is currently thought to be via rapid IFN-dependent reinforcement of antiviral intrinsic immunity throughout the body, to promote early viral control and host resistance upon primary acute infections^6, 7^. Yet, robust experimental data supporting this hypothesis are scarce, both in humans and mice. Indeed, unlike patients who do not respond to IFN-I/III^18, 19^, patients with homozygous loss-of-function mutations in TLR7, MYD88, IRAK-4 or IRF7 do not show increased susceptibility to most viral infections, with the exception of Influenza or SARS-Cov2^11, 12, 13, 14, 15, 16, 17^. However, the role of loss of pDC IFN production in this increased susceptibility to respiratory viral infections is unclear, since other cell types and biological processes are also directly affected by the same mutations. In mice, although pDCs are a major IFN sources during many viral infections, their depletion or functional inactivation did not compromise host survival, except upon systemic infection with HSV-2 or ocular infection with HSV-1 ^61, 62^. Here, we showed that during systemic infection with MCMV, a natural rodent pathogen, pDCs were the main source of both IFN-Is and IFN-IIIs, consistent with previous studies^30, 35^. However, the selective ablation of pDCs compromised neither the IFN-dependent induction of ISGs, nor viral control or host survival. These results are consistent with our previous work showing a greater impact of *Ifnar1* inactivation over *Myd88* deficiency on ISG expression and host survival in MCMV-infected mice^20^. Moreover, we also showed that ISGs were induced early in the lungs of mice infected with Influenza or SARS-CoV2, irrespective of the presence of pDCs, and that pDCs were not required for resistance to these infections. This was also true in mice expressing a functional allele of *Mx1*, the key restriction factor against IAV. Hence, the increase in IAV titers observed previously in *Mx1*^+^ mice depleted with anti-Bst2 antibodies ^43^ was unlikely to have been caused by pDC depletion but rather resulted from the perturbation of other cell types also targeted by this treatment. Thus, contrary to the prevailing dogma in the field, our results demonstrate that pDCs are redundant with other cellular sources of IFN for enhancing antiviral intrinsic immunity and promoting overall host resistance to several systemic or respiratory viral infections in mice.

Although immune cell activation was impaired in the absence of pDCs in several models of viral infections, including those used in our study^20, 30, 38, 61, 63^, these defects were transient and had very limited impact on host resistance to infection, suggesting the existence of other redundant mechanisms able to preserve the induction of protective innate and adaptive antiviral immune responses in the absence of pDC functions. Our findings thus emphasize the need to reevaluate whether and how pDC functions could benefit the host during viral infections. The use of pDC-less mice and of shield BM chimeras derived thereof will certainly be key to address these questions in future studies.

In the present study, during IAV or SARS-CoV2 infections of mice, we showed not only that pDCs were dispensable but also that they could actually be detrimental to host survival. Specifically, in K18:hACE infected mice, pDCs were dispensable in females, but deleterious in males. This appears contradictory with the prominent hypothesis in the field that pDCs are protective against human respiratory infections^11^. However, the picture is more complex as the role of pDCs in respiratory Influenza and SARS-CoV2 infections remains debated. In mouse models of Influenza infection, different studies have drawn contrasting conclusions, proposing that pDCs are beneficial^43^, dispensable^41, 42^ or deleterious^44, 45, 46^. However, these studies used experimental strategies to deplete pDCs or impair their functions that were not specific enough and also targeted other cell types or biological processes, potentially confounding the interpretation of the results in terms of the contribution of pDCs to the phenotypes observed. For example, previous studies showed a deleterious role during murine IAV infection of Bst2^+^ cells^44, 45, 46^, encompassing not only pDCs but also inflammatory monocytes, activated DCs and B cell subsets. In patients with severe COVID, blood pDCs were numerically reduced and defective in IFN production upon *ex vivo* stimulation^16, 64, 65, 66^. Moreover, an enhanced frequency of severe COVID19 was detected in patients with loss-of-function mutations disrupting the TLR7-to-MYD88-to-IRF7 signaling pathway that is essential for pDC IFN production. These observations led to the proposal that pDCs are protective against COVID in most patients and that disruption of their responses contribute to disease severity. However, alternative interpretations should be considered. The decrease in pDC numbers and the loss of their ability to produce IFN in the blood of severe COVID patients could reflect the recruitment and activation of these cells into the infected tissues where they could contribute to fuel inflammation by producing IFN locally^67^, as it has been reported to occur during infections of macaques or humans by immunodeficiency viruses^68, 69^. The protective role of the TLR7-MYD88-IRF7 signaling pathway observed in human respiratory viral infections could also occur in cells other than pDCs, in particular in monocytes or macrophages, which also express, and respond to, these molecules^16, 70, 71^. Indeed, other studies proposed that pDCs are a deleterious source of IFNs promoting exacerbated activation of monocytes/macrophages responsible of life-threatening immunopathology ^51, 52, 53, 54^. Here, we showed that, during SARS-CoV2 infection of mice, pDCs can be dispensable or deleterious, whereas Tlr7 and IFN-I responses are protective. pDCs are increased in the lungs and contribute to local IFN-α production during infection. However, Tlr7 deficiency more severely and broadly decreases both IFN-α and IFN-β production in the lungs. Thus, our findings support a scenario where the IFN-Is produced by pDCs are not required for survival and can even be detrimental, whereas other Tlr7-dependent cellular sources of IFN-I are critical for host resistance. Future studies are required to test this hypothesis, especially to compare the kinetics and spectrum of the TLR7-dependent production of IFN subtypes between pDCs and other cells *in vivo* during SARS-CoV2 infection. These experiments could be performed in mice, but also potentially in macaques, as the animal model naturally susceptible to this infection that the most closely mimics human disease. This has important implications as it has been proposed to boost pDC IFN production to treat severe human respiratory infections^8^, which could lead to an outcome exactly opposite to expectations if pDCs are deleterious in these diseases as we propose. On the contrary, understanding whether and how pDC IFN production could worsen disease may uniquely allow to disentangle the mechanisms underlying the beneficial versus deleterious effects of IFN responses in these diseases. This could ultimately allow designing intervention strategies to inhibit the deleterious effects of IFNs while still preserving their beneficial functions.

Most IFN-α subtypes, except IFN-α4 and IFN-β, were reduced in the lungs of IAV-infected pDC-less mice. The production of most IFN-α subtypes requires IRF7 that is constitutively expressed in pDCs but not in most other cell types, whereas IFN-α4/β can be quickly released by IAV-infected cells in an IRF3-dependent IRF7-independent manner^72, 73^. Although all IFN-Is interact with the same IFNAR receptor complex, IFN-I subtypes differ in their affinity and duration of binding to IFNAR, leading to differences in the intensity, kinetics and array of ISG induction. In addition, differently from most infected cells, pDCs are highly motile, especially after activation^63^, thus being able to release their IFNs to specific cells in a precise time and location, as we previously showed during MCMV infection^32, 37^. Hence, a division of labor may exist between different types of IFN-producing cells. Virally-infected cells could be mainly involved in rapidly reinforcing intrinsic immunity locally, in all neighboring cells, to control viral propagation. As a complementary response, pDCs could travel within infected tissues and across organs to deliver IFNs at slightly later time points to conventional DCs or other immune cell types to exert unique immunostimulatory functions. This pDC response may be generally beneficial but could become detrimental during infections that trigger IFN-dependent immunopathology or autoimmunity.

In conclusion, contrary to the dogma currently prevailing in the field, we show that, in mice, pDCs appear to be a largely dispensable source of IFN-I for host resistance to primary acute viral infections, similarly to what had been proposed several years ago in humans^9, 10, 74^. These findings illustrate how redundancy contributes to the robustness of immune responses ^20, 75^, including the signaling pathways and cellular sources leading to the production of IFNs during viral infections which is vital for the host^76^. However, our results also raise the question of the identification of the exact benefit for the host of the evolutionary conservation of pDCs, especially considering that current state of knowledge supports their detrimental roles in both infection-induced and sterile inflammation, whereas rigorous experimental proof are scarce of any beneficial role of pDCs in the natural history of diseases. Several hypotheses can be formulated. First, pDCs might be required for the protection of specific immune-privileged organs that are otherwise unable to produce protective IFNs, as recently shown for the eye in a mouse model of cornel infection with HSV-1^62^. Second, rather than being required for resistance against a primary acute infection with a first virus, pDCs may be required to induce a body-wide state of antiviral resistance including in barrier tissues remote from the initial site of infection^77^, in order to prevent superinfections by other viruses that could otherwise constitute a second lethal hit for the host. Third, pDC IFN production during a first primary acute viral infection could improve the induction of adaptive immune memory to enhance host protection to secondary infections, explaining in part the enhanced immune resistance of mice exposed to a normalized microbial environment^78^. Further studies will be necessary to investigate these hypotheses, which will require the generation of a novel mouse model of conditional rather than constitutive pDC depletion, not suffering from the confounding side effects reported for existing models^31, 35^, and complex experimental set-up of sequential exposure to different pathogens^79^.

## Methods

### Mice

All animal experiments were performed in accordance with national and international laws for laboratory animal welfare and experimentation (EEC Council Directive 2010/63/EU, September 2010). Protocols were approved by the Marseille Ethical Committee for Animal Experimentation (registered by the Comité National de Réflexion Ethique sur l’Expérimentation Animale under no. 14; APAFIS no. 21626-2019072606014177 v.4 and APAFIS #35098-2022020215455814 v4, APAFIS#26484-2020062213431976 v6). C57BL/6J (B6) mice were purchased from Janvier Labs. Siglech^iCre^ (*B6-Siglech^tm1(iCre)Ciphe^*), Ifnar1-KO (B6.129S2-Ifnar^1tm1agt^) and SCRIPT (Siglech^iCre^;Pacsin1^LSL-tdT^;*Ifnb*^EYFP^) mice were previously described^32,48^. *Pacsin1*^LoxP-STOP-LoxP-DTA^ (*B6-Pacsin1^tm1(DTA)Ciphe^*, Pacsin^LSL-DTA^) and Stat1-KO (B6-Stat1^tm1d(EUCOMM)Ciphe^) were generated by CIPHE. Homozygous Pacsin1^LSL-DTA^ and Siglech^iCre^ were bred together and their double heterozygous Siglech^iCre^; Pacsin1^LSL-DTA^ progeny was named pDC-less mice. Siglech^iCre^ and Pacsin^LSL-DTA^ were also bred with CD45.1 congenic mice, then interbred to generate 5.1 pDC-less mice used as recipient for Shield Bone Marrow Chimera (SMBC) generation. TLR7^KO^ (*B6-Tlr7^tm1Flv^)* were kindly provided by L. Alexopoulou, CIML. B6.A2G-*Mx1^+/+^* congenic mice were generated and provided by M. Le Bert, TAAM, Orleans, France. IR7^KO^ (*B6-Irf7^tm1Ttg^)* were kindly provided by C. Svanborg, Lund, Sweden. *Ifnb*^EYFP^ (B6.129-Ifnb1^tm1Lky^), hBDCA2-DTR (C57BL/6-Tg (CLEC4C-HBEGF)956Cln/J) and heterozygous K18-hACE C57BL/6J mice (strain: 2B6. Cg-Tg (K18-ACE2)2Prlmn/J) were obtained from Jackson Laboratories, USA. Pacsin^LSL-DTA^ and B6.A2G-*Mx1*^+/+^ mice were intercrossed to generate double homozygous mice, then bred with Siglech^iCre^ to obtain as progeny *Mx1*^+^ pDC-less mice. Control mice used in all experiments were Pacsin^LSL-DTA^, Siglech^iCre^ or B6 mice. All mouse strains were on B6 genetic background and bred at the Centre d’ImmunoPhénomique (CIPHE) or the Centre d’Immunologie de Marseille-Luminy (CIML), under specific pathogen free-conditions and in accordance with animal care and use regulations. Mice were housed under a 12 h dark:12 h light cycle, with a temperature range of 20–22 °C and a humidity range of 40–70%. All animals used were sex and age matched (8-12 weeks of age for all experiments, except for SBMC). C57BL/6, coming from an external provider, as well as Siglech^iCre^ and Pacsin^LSL-DTA^ mice, bred in the same mouse house as pDC-less mice, gave similar readouts and were indistinctly used as control in experiments with MCMV or IAV.

### Shield Bone Marrow Chimera (SMBC) generation

5 weeks-old CD45.1+ pDC-less were anesthetized, then their hind legs were 9 Gy irradiated, while the rest of the body was protected with a lead shield. At the end of the irradiation, mice were intravenously injected with 15 x 10^6 cells of BM isolated from indicated CD45.2+ donor mice. 4 weeks after BM engraft mice were bled to test the reconstitution with donor BM cells, then used for experimentation.

### Viruses, viral infections and mice treatment

MCMV Smith stocks were prepared from salivary gland extracts of 3-week-old, MCMV-infected BALB/c mice. Mice were infected intraperitoneally with 10^5^ Plaque-Forming Units (PFU) and sacrificed at the indicated time points. For survival experiments with MCMV, mice were infected with 1.4 x 10^5^ PFU. H3N2 A/Scotland/20/74 IAV strain was produced and titrated in vitro by using Madin-Darby Canine Kidney (MDCK) cell line. Mice were anesthetized with ketamine/xylazine, then intranasally infected with 40µl of DMEM medium containing 160 PFU of IAV strain and euthanized at indicated time points. For survival studies mice were infected with 145 PFU of the same IAV strain. β CoV/France/IDF0372/2020 SARS-Cov2 strain was supplied by the National Reference Centre for Respiratory Viruses hosted by the Institut Pasteur (Paris, France). This strain was isolated from a human sample provided by the Bichat Hospital, Paris, France and corresponded to the original Wuhan/D614 SARS-Cov2 strain detected at the beginning of COVID19 pandemic. Infectious stocks were grown by inoculating Vero E6 cells and collecting supernatants upon observation of the cytopathic effect. Debris was removed by centrifugation and passage through a 0.22 mm filter. Supernatants were stored at -80 °C. Mice were anesthetized with 150μL of ketamine/xylazine, then intranasally infected with 30µl of medium containing 200 PFU of β CoV/France/IDF0372/2020 SARS-CoV2. For Bulk RNASeq studies mice were infected with a high dose (1.1 x10^5^ PFU) of SARS-CoV2 strain. IAV-infected and SARS-CoV2 infected mice were monitored daily for morbidity (body weight) and mortality (survival). During the monitoring period, mice were scored for clinical symptoms (weight loss, eye closure, appearance of the fur, posture, and respiration). Mice obtaining a clinical score defined as reaching the experimental end-point were humanely euthanized, according to experimental protocols approved by ethical committee. For pDC depletion in hBDCA2-DTR mice, mice were intraperitoneally injected with 15ng/g of Dyphtheria Toxin, Merck, at day 2 after IAV infection. For neutrophil depletion mice were intraperitoneally injected at day 5 and 6 after IAV infection with 500 µg/mouse of anti-Ly6G (1A8) or isotype control antibodies, both purchased from Biolegend.

### Cell preparation

Mice were euthanized and perfused with PBS 1×. Spleen and lymph nodes were digested for 25 min at 37 °C with Collagenase IV (Worthington biochemical) and DNase I (Roche Diagnostics). Organs were then mechanically crushed and filtered over 70-µm cell strainers (Corning). Bone marrow cells were flushed from mouse femurs. Livers and lungs were harvested, minced and mechanically digested with Collagenase IV and DNase I by using GentleMACS (Miltenyi, program 37_m_LIDK_1 for the liver, 37_m_LDK_1 for the lungs). At the end of the digestion, lung cell suspensions were filtered over 70-µm cell strainers and cells were pelleted by centrifugation. Liver cells were submitted to a 80:40 Percoll gradient. Small intestines were harvested, opened longitudinally, cut into 1-cm pieces and washed extensively with PBS 1×, then incubated 3× at 37 °C on shaking (200 r.p.m.) with PBS 1× containing 2% fetal calf serum (FCS) and 5 mM EDTA. At the end of each incubation, supernatants were collected and centrifuged. Pelleted cells from the three incubations were pooled together and submitted to a 67:44 Percoll gradient. Bronchoalveolar lavages (BAL) were performed upon intratracheal injection of 1 ml of cold PBS, repeated once. BAL were then centrifuged, fluids harvested and frozen at -80°C until their use. RBC lysis was performed with RBC lysis buffer (Thermofisher). For the lungs, the left lobe was harvested for RNA extraction, while the other lobes were used for flow cytometry.

### Flow cytometry analysis

Cells were first incubated with 2.4G2 mAb for 10 minutes at 4°C, then we performed extracellular staining in staining buffer, PBS 1× supplemented with 2 mM EDTA (Sigma-Aldrich) and 2% FCS for 30 min at 4 °C. Dead cell staining (LIVE/DEAD Fixable Aqua Dead Cell Stain, Life Technologies) was performed in PBS 1× according to the manufacturer’s recommendations. All antibodies used and their related dilution have been listed in the Reporting Summary of this manuscript. Before acquisition samples were fixed with Cytofix/Cytoperm BD 1X (eBioscience) or with a 2% formaldehyde solution in PBS 1× when using cells expressing EYFP protein. Samples were acquired with a FACS LSR UV (BD Biosciences) using BD Diva v.9.0. All data were analyzed with FlowJo v.10.8.1 software.

### RNA extraction and RT-qPCR

Harvested organs were incubated in RNAlater (Thermofisher) at 4°C overnight. Total RNAs from lung cells were extracted using RNeasy Plus Mini Kit (Qiagen) following the manufacturer protocol. Retrotranscription into complementary DNA (cDNA) was performed using the Quantitect Reverse Transcription Kit (Qiagen). The expression levels of the following murine genes were determined by quantitative PCR (qPCR) using the SYBR® Premix Ex TaqTM kit and analysed using the Prism 7500 Fast PCR System. Relative gene expression was calculated using the ΔΔCt method with *Actinb* as housekeeping gene for normalization. The primers used were as it follows: *Actinb* forward 5’-GGCTGTATTCCCCTCCATCG-3’; reverse 5’-CCAGTTGGTAACAATGCCATGT-3’; *Il28b* forward 5’— GGAGGCCCAGAGCAAGGA-3’ ; reverse 5’-TTGAAACAGGTTGGAGGTGACA-3’ ; *Irf7*forward 5’— CCACGCTATACCATCTACCTGG-3’ ; reverse 5’-GCTGCTATCCAGGGAAGACAC-3’ ; *Isg15* forward 5’— GGTGTCCGTGACTAACTCCAT-3’ ; reverse 5’-TGGAAAGGGTAAGACCGTCCT-3’ ; *Mx2* forward 5’— AGAGGGAGAATGTCGCCTATT-3’ ; reverse 5’-CGTCCACGGTACTGCTTTTCA-3’; *Rsad2* forward 5’— TGCTGGCTGAGAATAGCATTAGG-3’ ; reverse 5’-GCTGAGTGCTGTTCCCATCT-3’ ; *Ifna1* forward 5’-GGATGTGACCTTCCTCAGACTC-3’; reverse 5’- ACCTTCTCCTGCGGGAATCCAA-3’ ; *Ifna4* forward 5’— AAGGACAGGAAGGATTTTGGAT-3’ ; reverse 5’-GAGCCTTCTGGATCTGTTGGTT-3’ ; *Ifna5* forward 5’- GGATGTGACCTTCCTCAGACTC-3’; reverse 5’-CACCTTCTCCTGTGGGAATCCA-3’ ; *Ifnb1* forward 5’-GGTGGTCCGAGCAGAGATCTT-3’; reverse 5’-CAGTTTTGGAAGTTTCTGGTAA-3’.Viral titration of MCMV was performed as previously described^20^. For viral titration of IAV, we performed RT-qPCR by testing the expression of *M1* IAV gene, using the following primers: forward 5’—AAGACCAATCCTGTCACCTCTGA-3’; reverse 5’-CAAAGCGTCTACGCTGCAGTCC-3’. For viral titration of SARS-CoV2, tissues were homogenized with ceramic beads in a tissue homogenizer (Precellys, Bertin Instruments) in 0.5 mL RLT buffer, or lung tissues were harvested, minced and mechanically digested with Collagenase IV and DNase I by using GentleMACS (Miltenyi, program 37_m_LIDK_1 for the liver, 37_m_LDK_1 for the lungs). At the end of the digestion, lung cell suspensions were processed with RTL buffer. RNA was extracted using the RNeasy Mini Kit (QIAGEN) or Direct-zol RNA Mini PrepPlus de ZYMO RESEARCH, and reverse transcribed using the High-Capacity cDNA Reverse Transcription Kit (Thermo Fisher Scientific). Amplification was carried out using OneGreen Fast qPCR Premix (OZYME) according to the manufacturer’s recommendations. Copy numbers of the RNA-dependent RNA polymerase (*RdRp*) SARS-CoV2 gene were determined using the following primers: forward 5’-CATGTGTGGCGGTTCACTAT-3’, reverse 5’-GTTGTGGCATCTCCTGATGA-3’. This region was included in a cDNA standard to determine the copy number determination down to 100 copies/reaction. SARS-CoV-2 copy numbers were compared and quantified using a standard curve and normalized to total RNA levels. An external control (mock-infected wild type animal) and a positive control (SARS-CoV2 cDNA containing the targeted region of the *RdRp* gene at a concentration of 10^4^ copies/µl [1.94x10^4^ copies/µl detected in the assay]) were used in the RT-qPCR analysis to validate the assay.

### Bulk RNA-Seq

RNA was isolated from the lungs of uninfected and IAV-infected control and pDC-less mice (160 PFU, days 0, 3 and 6 after infection) or of uninfected and SARS-CoV2 infected K18:hACE2 mice (1.1 10^5 PFU, days 0, 1, 2, 4 and 6 after infection), as described in RNA extraction method. For Bulk RNASeq 500 ng of lung RNA/sample were used to prepare libraries for sequencing by using KAPA RNA HyperPrep Kits (Roche). Libraries generated with IAV-infected samples were sequenced by Macrogen, while libraries generated from SARS-CoV2 infected samples were sequenced at the Genomic facility of the CIML. All libraries were sequenced using Illumina platforms. Alignments of fastq files obtained after sequencing were generated with STAR 2.7.9a software using the GRCm38 reference genome. The number of reads mapped to each gene was determined with featureCounts v2.0.1 using Mus_musculus.GRCm38.100.gtf annotation file. Gene models (gene symbols of the type “Gmxxxx”) were then removed from the feature count file and TPM normalization was applied using the Scuttle package of R 1.0.34.

### FITC-Dextran

To assess lung permeability, isoflurane-anesthetized mice were administered intratracheally with 10 μg/mouse (50µl of a solution at 200µg/ml in PBS 1x) of FITC-dextran (Sigma) 8 days after IAV infection. 1 hour later, blood was collected from anesthetized mice from the retro-orbital sinus and the plasma was separated by centrifugation. Plasma were diluted 1:2 in PBS 1X. Dextran leakage in the bloodstream was measured as FITC fluorescence in the plasma. Fluorescence was read at 520 nm with excitation at 490 nm by using a Mithras LB 940 plate reader (Berthold Technologies).

### Cytokine quantification

Cytokines and chemokines present in the sera of uninfected vs MCMV-infected mice and in the BAL of uninfected vs IAV-infected mice were analyzed using the LEGENDplex Mouse Anti-Virus Response Panel (BioLegend). Samples and standards were plated in technical duplicates and the assay was executed according to the manufacturer’s protocol. Data were collected using the CANTO II (BD Biosciences) and analyzed using the cloud-specific LEGENDplex Data Analysis Software Qognit (BioLegend).

### Histological analysis

Lungs were harvested from control and pDC-less mice 8 days after IAV infection and kept with 10% buffered formalin (VWR Chemical) for 24 hours, then dehydrated and embedded in paraffin (Fischer Histoplast). 3,5µm-thick sections were cut using the microtome Leica RM2245. Hematoxylin-eosin staining was made automatically with Leica Autostainer XL. Finally, the slides were mounted with Entellan mounting medium (Merck) and kept at room temperature. Tissue sections were blindly assessed by a certified veterinary pathologist. Microscopic examination was done using a Nikon Eclipse E400 microscope. Pictures were taken with a Nikon DS-Fi2 camera and NIS Elements imaging software (Nikon, Japan). Histopathological semi-quantitative lung inflammation scoring system was performed according to Supplementary Table 1.

### Immunohistofluorescence, confocal microscopy and image analysis

Mouse lungs were perfused with a mixture of 1:2 of AntigenFix (Diapath, containing 4% paraformaldehyde) and 1:2 of Optimal Cutting Temperature (OCT, Sakura Fineteck), collected, fixed in AntigenFix for 4h at 4°C and then washed several times in phosphate buffer (PB; 0.025 M NaH2PO4 and 0.1 M Na2HPO4). Lungs were incubated in PB solution containing 30% Sucrose overnight at 4°C and then embedded in OCT freezing medium, snap frozen and stored at −80 °C. 30 μm-thick cryosections were performed using a microtome (Leica 3050s Cryostat) at temperatures between −24 °C and −20 °C.

For immunostaining, lung sections were blocked with PB, 0.1% Triton X-100 and 2% BSA for 1h at room temperature and then stained overnight at 4°C with primary antibodies diluted in PB, 0.1% Triton X-100 and 2% BSA. After several washings with PB, lung sections were then stained in PB, 0.1% Triton X-100 and 2% BSA with secondary antibodies for 2h at 4°C. To detect YFP signal, after the incubation with secondary antibodies, sections were washed and incubated with PB, 0.1% Triton X-100, 2% BSA and 5% rabbit serum for 45 min at room temperature. Sections were then stained with Alexa 488-conjugated anti-GFP antibodies for 1h at 4°C. Finally, lung sections were washed with PB and mounted with a coverslip and Prolong Antifade Gold mounting medium (Life Technologies). Stained lung sections were acquired using spectral confocal microscopes (Zeiss LSM880 or Zeiss LSM980) with x10/0.45 or x20/0.8 objectives, respectively. Images were acquired using the ZEN blue software (Carl Zeiss Microscopy), in 16-bit format, in spectral mode to suppress autofluorescence for each fluorochrome used. Images were then analyzed using ImageJ software ^80^. For CD45 quantification areas expressing low versus high CD45 signal were qualitatively identified in each whole lung section, then a region of interest (ROI) was created for each area and CD45 mean fluorescence intensity (MFI) was measured. In highly inflamed areas CD45 MFI was at least twice of that detected in the lowly inflamed areas. We then used QuPath v0.4.3 ^81^ for cell quantifications. CD45+ cells, IAV-infected HA+ cells and Tom+ pDCs were quantified using the QuPath cell detection module. Then, using the QuPath classifier module, we quantified among the Tom+ pDCs, the IFN+ (Tom+ YFP+) pDCs expressing at least a threshold value of 2216 in YFP channel. For lung integrity analysis, EpCAM+ areas were defined as areas expressing at least a 5000 threshold for EpCAM channel, then their percentage was calculated on the whole lung surface analyzed. EpCAM+ bronchi were classified as closed when EpCAM staining was continuous and as opened when EpCAM was disrupted. Both types of EpCAM+ bronchi were manually quantified and corresponding percentages were calculated.

### Statistical analysis

All statistical analyses were performed using GraphPad Prism 10 software. All results are expressed as mean ± s.e.m. All quantifications were performed with awareness of experimental groups, meaning not in a blinded fashion. Animals were matched in age and gender between experimental groups and each cage was randomly assigned to a treatment group. No animals or data were excluded under any circumstances. Statistical parameters including the exact value of n (number of biological replicates and total number of experiments) and the types of statistical tests are reported in corresponding legends. Comparisons between two groups were assessed using unpaired and non-parametric Mann–Whitney test. Multiple comparisons were assessed using a Kruskal-Wallis test followed by Dunn’s post-hoc test. Survival curves were analyzed using Log-rank (Mantel-Cox) test. Comparisons between groups were planned before statistical testing and target effect sizes were not predetermined. Each dot indicates an individual. p values have been specified in each panel.

## Acknowledgements

We thank CIPHE for the generation of *Pacsin1*^LSL-DTA^ mice and their assistance in the breeding of mice, as well as the staff of the CIML mouse houses, flow cytometry, histology, genomics, bioinformatics (CB2M) and imaging (ImagImm) core facilities. We thank Ramazan Akyol for his assistance during the generation of SBMC and Lionel Spinelli for his assistance for statistical analyses. We thank the Centre de Calcul Intensif d’Aix-Marseille for granting access to its high-performance computing resources. We thank Catharina Svanborg, Lund University, Sweden, for providing *Irf7*^KO^ mice. We thank Marie-Laure Dessain, TAAM, Orleans, France, for genotyping *Mx1* mice. This research was funded by the “Fondation pour la Recherche Médicale” (grant DEQ20180339172, Equipe Labellisée, to M.D.) and from the French National Research Agency (ANR, no. ANR-21-CO12-0001-01, ‘RIPCOV’ to M.D. and B.M., and no. ANR-23-CE15-0001, ‘RIPIREVI’, to M.D. and A.Z.). The projects that contributed to this publication also received funding from the DCBIOL Labex (ANR-11-LABEX-0043, grant no. ANR-10-IDEX-0001-02 PSL*), the Excellence Initiative of Aix-Marseille Universite-A*Midex (grants no. ANR-11-IDEX-0001-02 and AMX-20-TRA-008), and France-BioImaging (grant no. ANR-10-INBS-04) and PHENOMIN-Infrastructure (no.ANR-10-INBS-07), funded by the French government’s «Investissements d’Avenir» program managed by the ANR. We also acknowledge institutional support from CNRS (French National Centre for Scientific Research), INSERM (French National Institute of Health and Medical Research) and Aix-Marseille Université. C.N. is supported by a fellowship from the French Ministry of Research and Higher Education.

## Author contributions

C.N. designed, performed and analyzed most of the experiments, and contributed to write the manuscript. C.P.M. performed experiments for manuscript revision; K.R. performed RT-qPCR experiments on MCMV- and IAV-infected samples. M.V. prepared the libraries used for Bulk RNA-Seq samples and contributed to perform the experiments on SARS-CoV2-infected samples. N.C. genotyped and stabilized mutant pDC-less mice. G.B. performed viral titration on MCMV-infected samples. M.Fab., S.S., S.M., C.G., S.M. and A.S. bred K18:hACE and related double/triple mutant mice and performed the experiments with SARS-CoV2-infected mice and samples. F.F. directed, designed and analysed the generation of *Pacsin1*^LSL-DTA^ mice. C.L. performed anatomopathological analysis of lung sections from IAV-infected mice. L.A., M.G., C.G., A.B. and S.W. provided advice and shared reagents, mutant mice and experimental protocols. R.R. and N.H. respectively supervised and performed experiments on the brains of IAV-infected mice. P.M. provided advice and technical support for Bulk RNA-Seq experiments. B.E. and T.P.V.M. performed the mapping and the first analysis of Bulk RNA-Seq data. M. Fal. contributed to the setting of the analysis of confocal microscopy images. L.C. performed paraffin-embedding, prepared and stained lung sections used for anatomopathological studies. H.T. genotyped and provided *Irf7*^KO^ mice. T.B. provided advice and shared experimental protocols. M.L.B. provided B6.A2G-*Mx1*^+/+^ mice. B.M. contributed to direct and fund the study. A.Z., M.D. and E.T. contributed to fund the study and directed it, designed experiments, performed some experiments, analysed data and wrote the manuscript.

---

**Extended Fig. 1:**
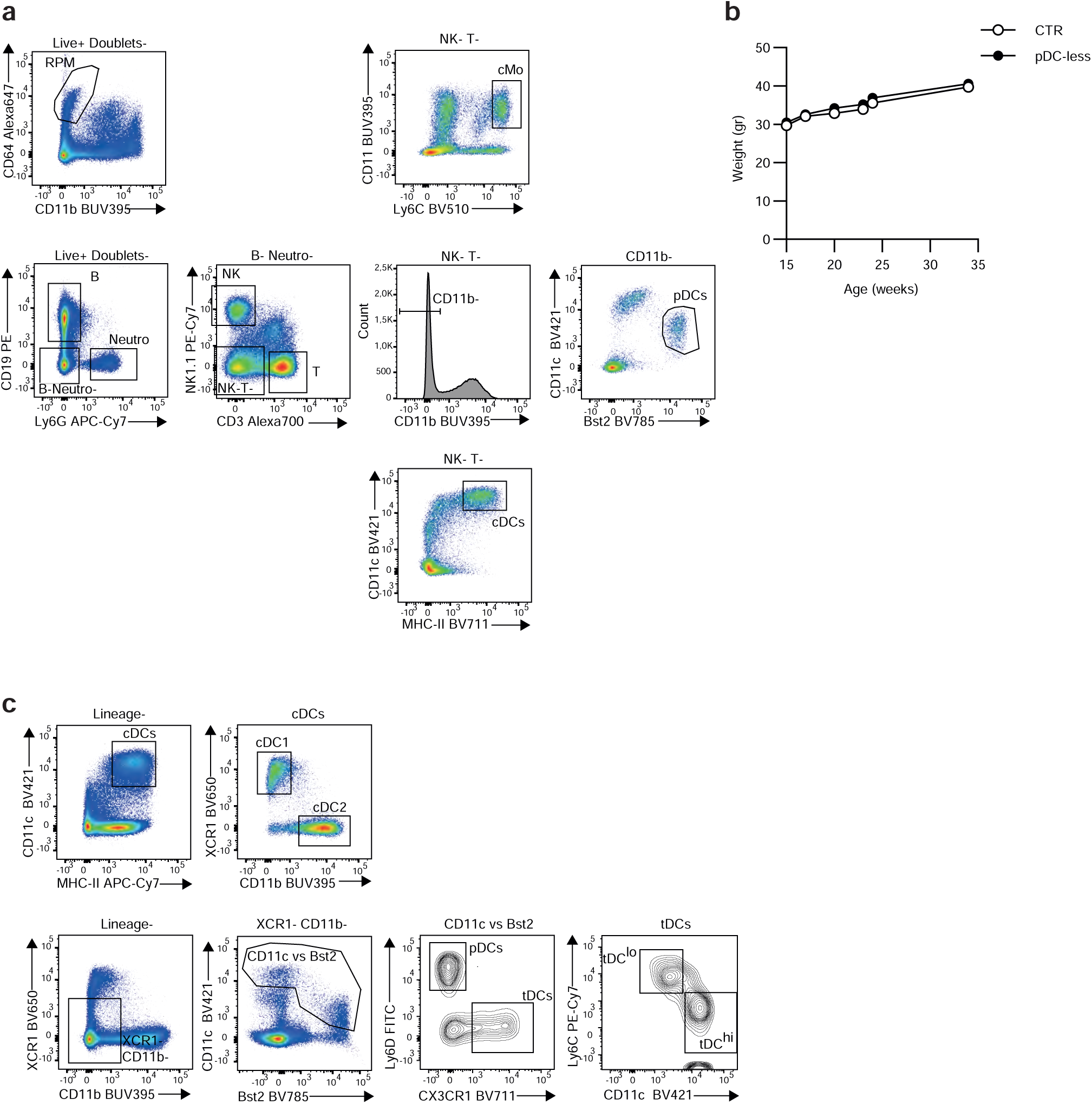
Gating strategy for the identification of myeloid and lymphoid lineages. a, c, Gating strategy used to identify myeloid and lymphoid immune cell types studied. **b**, The weight of control vs pDC-less was measured along time. Weight mean was represented at each time point analyzed for each strain of mice (n=5).

**Extended Fig. 2:**
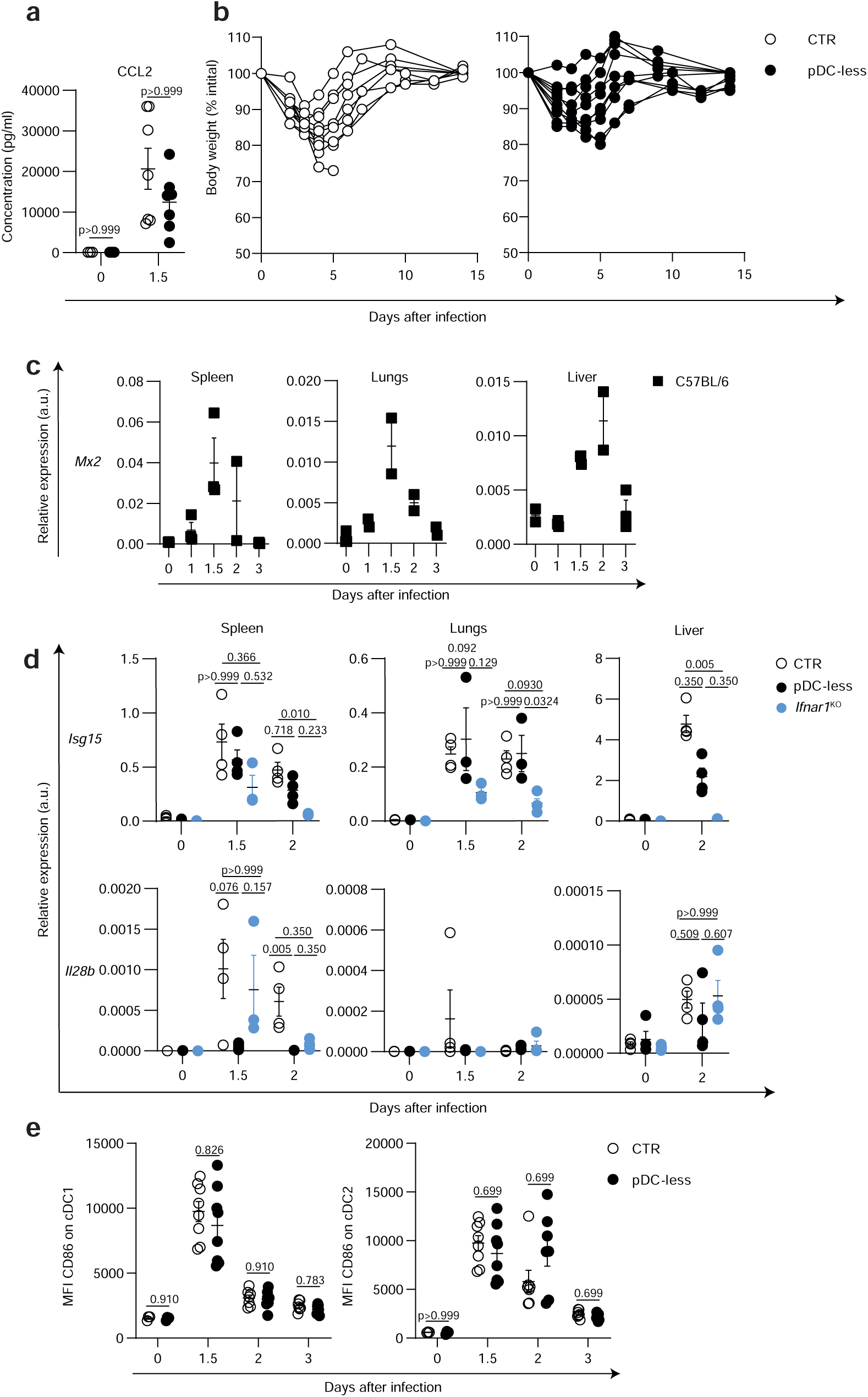
Complementary analysis of MCMV-infected control and pDC-less mice a, Cytometric bead arrays was performed to quantify CCL2 concentrations in sera isolated from uninfected (0) and 36h infected (1.5) control and pDC-less mice. Each dot represents an individual animal. The data shown (mean +/- s.e.m.) are pooled from two independent experiments (n=3 for 0, n=7 for 1.5). **b**, Weight curves of mice represented in Fig. 2a (10 CTR, 8 pDC-less). **c**, Relative expression of *Mx2* gene (mean +/- s.e.m.) was analyzed in the spleens of C57BL/6 infected with MCMV at indicated time points (n=3 for each time point). **d,** Expression levels of indicated genes was analyzed by RT-qPCR in indicated organs isolated from uninfected or MCMV-infected CTR, pDC-less and *Ifnar1*^KO^ mice at indicated days after infection. Data were normalized to expression level of *Actin* gene. The data (mean + s.e.m.) shown are from two independent experiments (n=4 for each condition at each time point). Statistical analysis was performed as described in Fig. 2d. **e**, CD86 MFI values on indicated cDC subsets (mean +/- s.e.m.) were determined in control and pDC-less mice at indicated days after MCMV infection (n=4 for 0, =8 for 1.5, =7 for 2 and =10 for 3 days after infection for both strains). Statistical analysis was performed as described in Fig. 2e.

**Extended Fig. 3:**
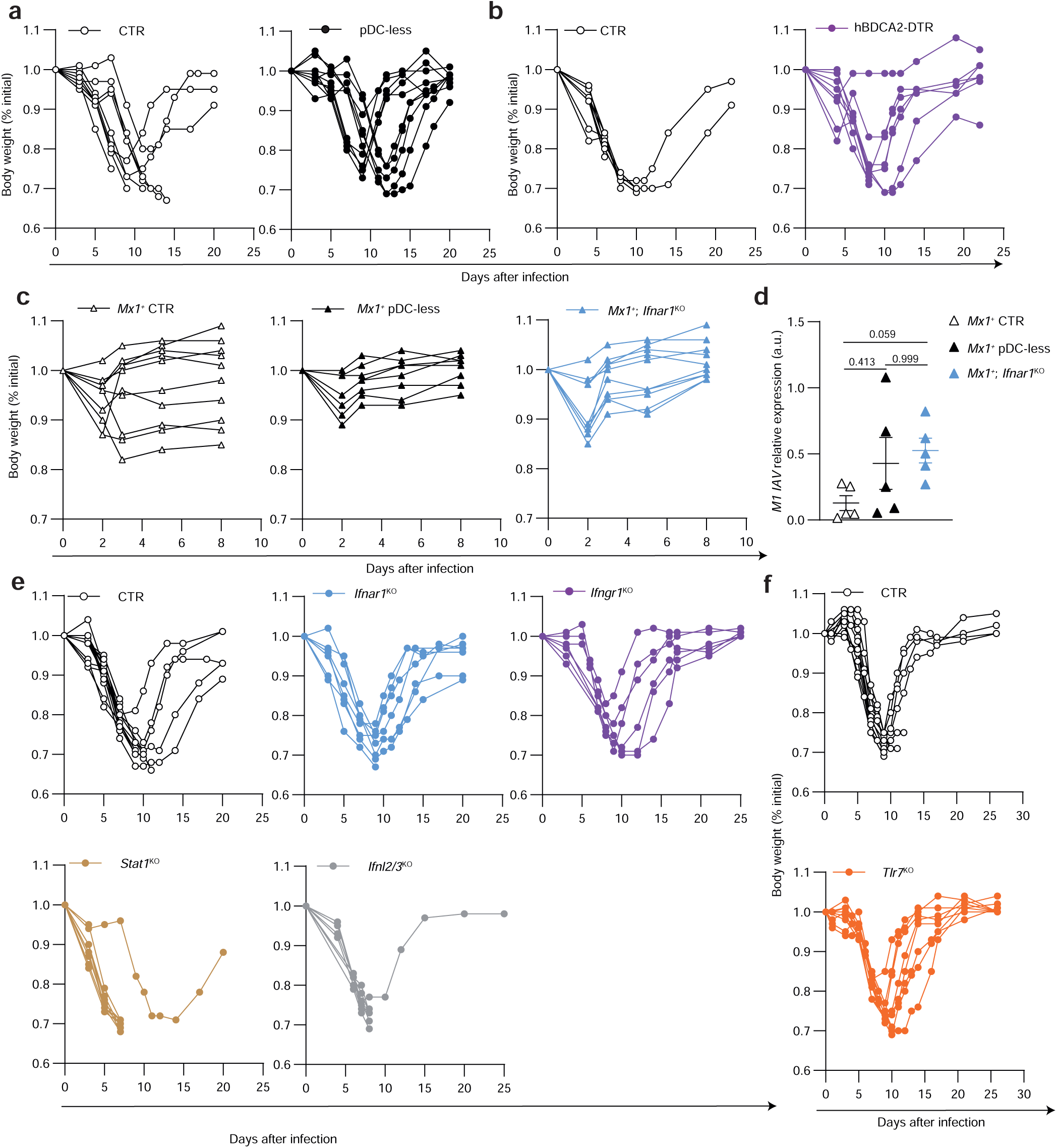
Weight curves of IAV-infected mice deficient in pDCs, in the response to IFN-I or to all IFNs, or in TLR7 a-c,e-f. The weight of indicated IAV-infected mice was regularly measured up to 2-3 weeks after infection. Data are represented as percentages of the weight at day 0 of the infection. Mice reaching the stop points were euthanized. **a**, n= 10 for both strains, **b**, =n=9 for both strains, **c**, n=10 *Mx1*^+^, 8 *Mx1*^+^ pDC-less, 10 *Mx1*^+^;Ifnar1^KO^ mice, **e**, n= 15 CTR, 8 *Ifnar1*^KO^, 7 *Ifngr1*^KO^, 9 *Ifnl2/3*^KO^ and 9 *Stat1*^KO^ mice. **f**, n= 10 CTR, 9 *Tlr7*^KO^ mice. **d**, M1 IAV expression was measured by RT-qPCR in the lungs of indicated mice 2 days after IAV infection. n=5 for each strain.

**Extended Fig. 4.**
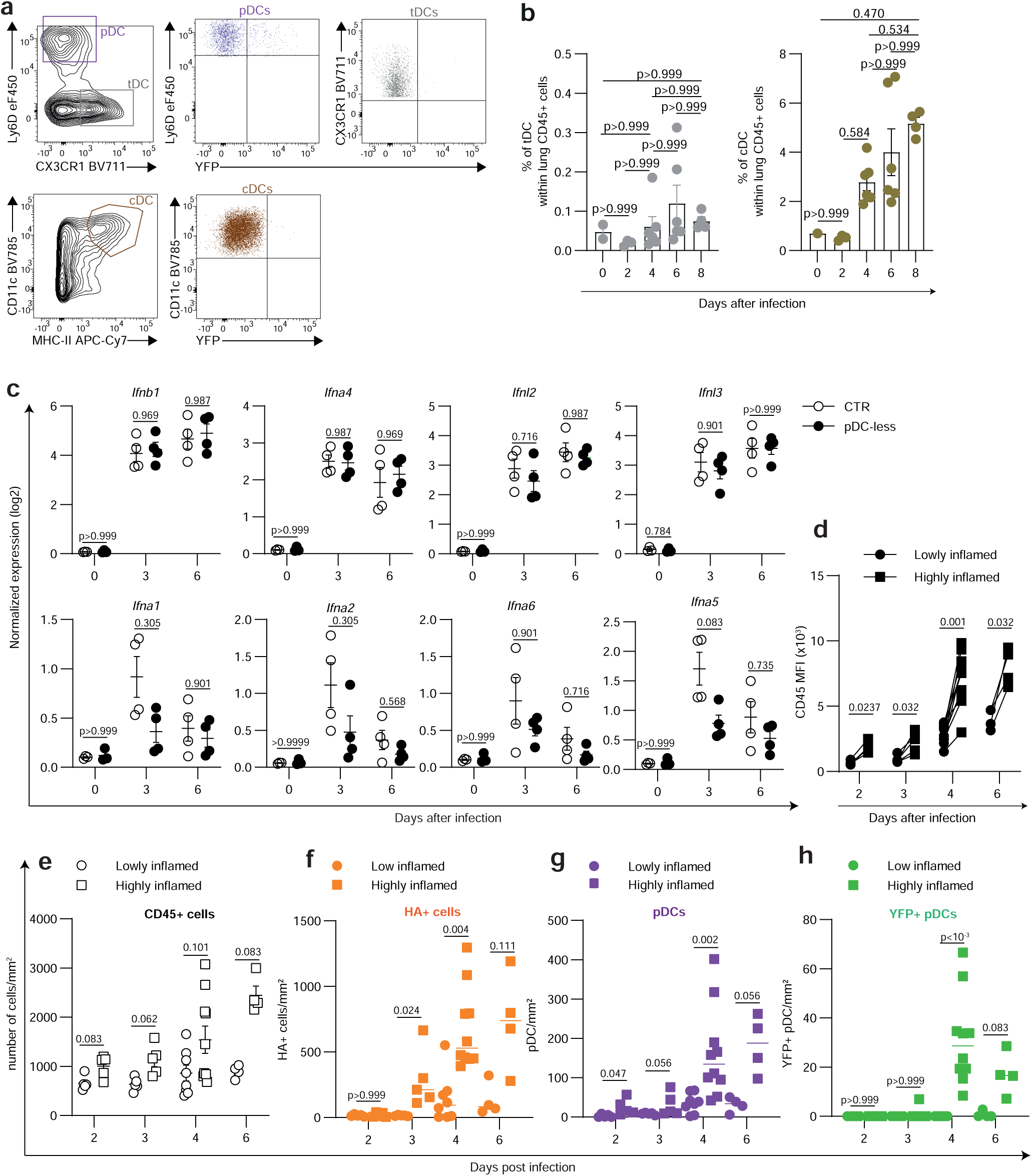
During IAV infection pDCs are increased in the lungs and are a main source of IFN-α. a-b, Gating strategy for (**a**) and percentages of (**b**) cDCs and tDCs isolated from the lungs of IAV-infected *Ifnb1*^EYFP^ mice. A two-tailed non-parametric Mann-Whitney test was used for the statistical analysis. n=2 for day 0, 3 for day 2, 6 for days 4 and 6, 4 for day 8 after infection. **c**, Normalized expression (log2) of indicated genes based on Bulk RNASeq samples obtained by whole lung samples isolated from CTR and pDC-less mice at day 0 (n=3/strain), 3 and 6 (n=4/strain) after IAV infection. **d**, Highly vs lowly inflamed areas were defined taking in account the CD45 MFI. For each tissue section CD45 MFI was at least twice in highly inflamed areas, as compared to lowly ones. **e-h**, The numbers of CD45+ cells (**e**), IAV+ cells (**f**), pDCs (**g**) and YFP+ pDCs (**h**) per mm² were quantified in highly and lowly inflamed areas in IAV-infected lungs. The data shown (mean +/- s.e.m.) in **d-h** are the pool of the analysis of two tissue sections/lung lobe/mouse for the following numbers of infected SCRIPT mice: n=5 at day 2, n=5 at day 3, n=10 at day 4 and n=4 at day 6 after infection. An unpaired and nonparametric multiple t-test (Mann-Whitney) with Holm-Sidak method for correction was used for the statistical analysis.

**Extended Fig. 5:**
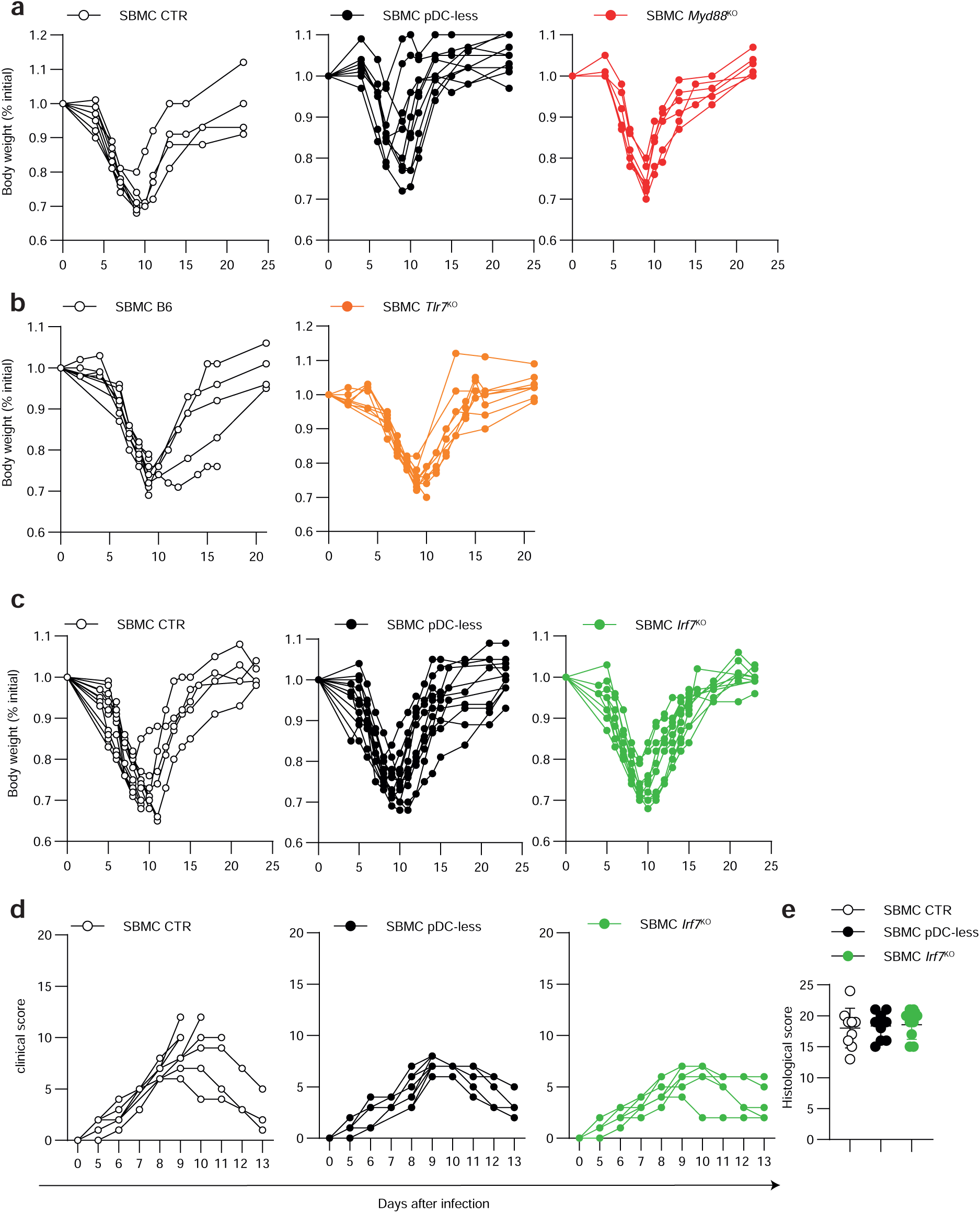
Weight curves of IAV-infected pDC SBMC mice deficient in pDCs or in molecules involved in Tlr7/Myd88/Irf7 signaling pathway. a-c, The weight of indicated IAV- infected mice was regularly measured up to 3-4 weeks after infection. Data are represented as percentages of the weight at day 0 of the infection. Mice reaching the end points were euthanized. **d**, Clinical scores of indicated IAV-infected SBMC were evaluated daily based on weight loss, eye closure, appearance of the fur, posture, and respiration. **a-b,** Pool of two independent experiments (n=8 for each strain); **c-d,** Pool of three independent experiments (n=12 for each strain). **e**, Histological score was performed on the lungs of indicated SBMC at day 8 after infection. n=10 for each pDC SBMC type.

**Extended Fig. 6:**
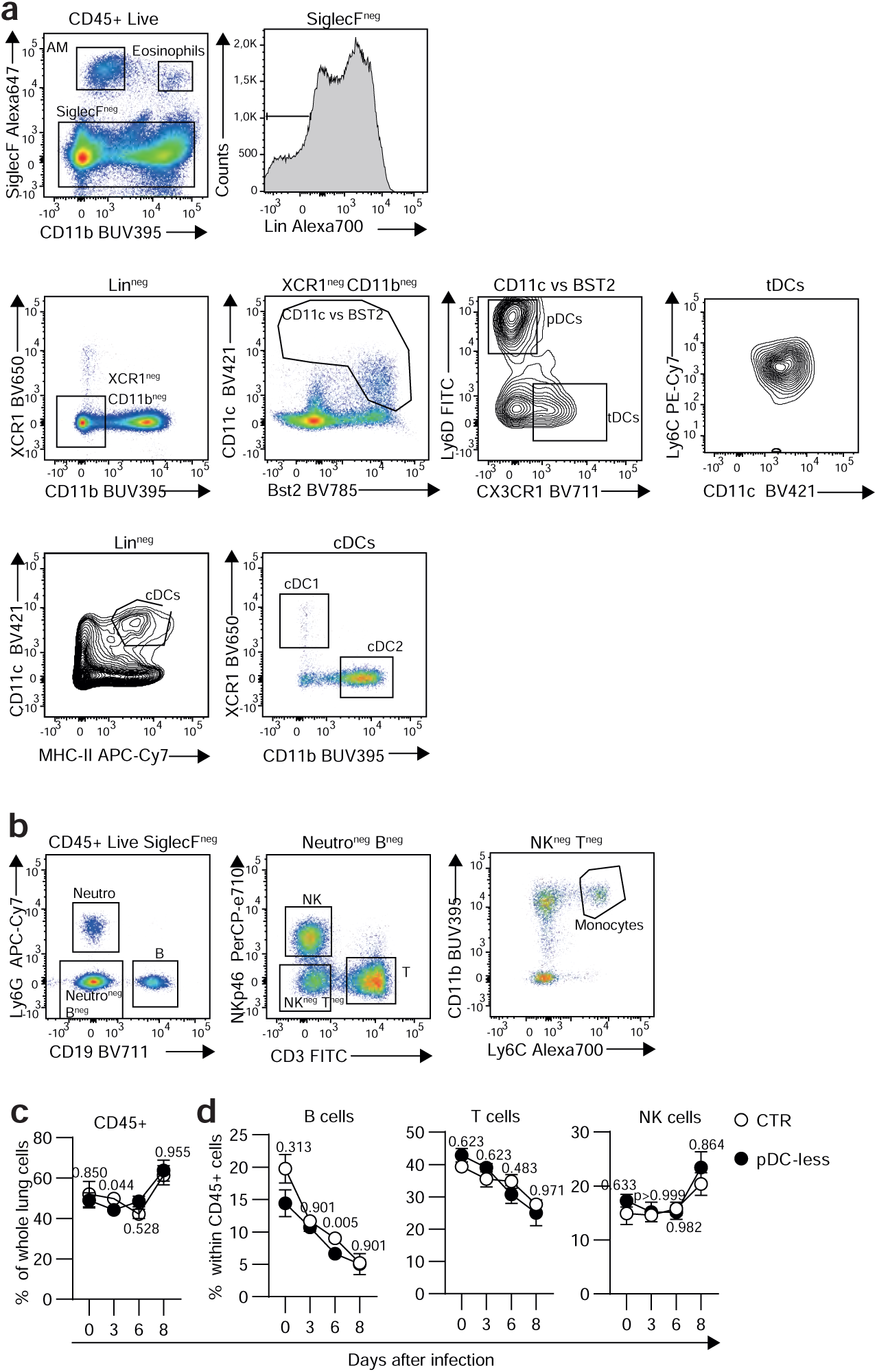
Gating strategy used to identify immune cell types in cells isolated from the lungs of uninfected vs IAV-infected control and pDC-less mice. a-b, Cells were isolated after enzymatic digestion of lungs of uninfected or IAV-infected animals and analyzed by flow cytometry. Gating strategy used to define indicated immune cell types has been depicted. Plots shown are from one representative mouse out of 40 analyzed. **c,d**, Percentages of CD45+ and lymphoid immune cells isolated from the lungs of control and pDC- less mice isolated at the indicated day after infection. For each mouse strain, n= 5 at day 0, 7 at day 3, 8 at day 6 and 4 at day 8. An unpaired and nonparametric multiple t-test (Mann-Whitney) with Holm-Sidak correction method was used for the statistical analysis.

**Extended Fig. 7:**
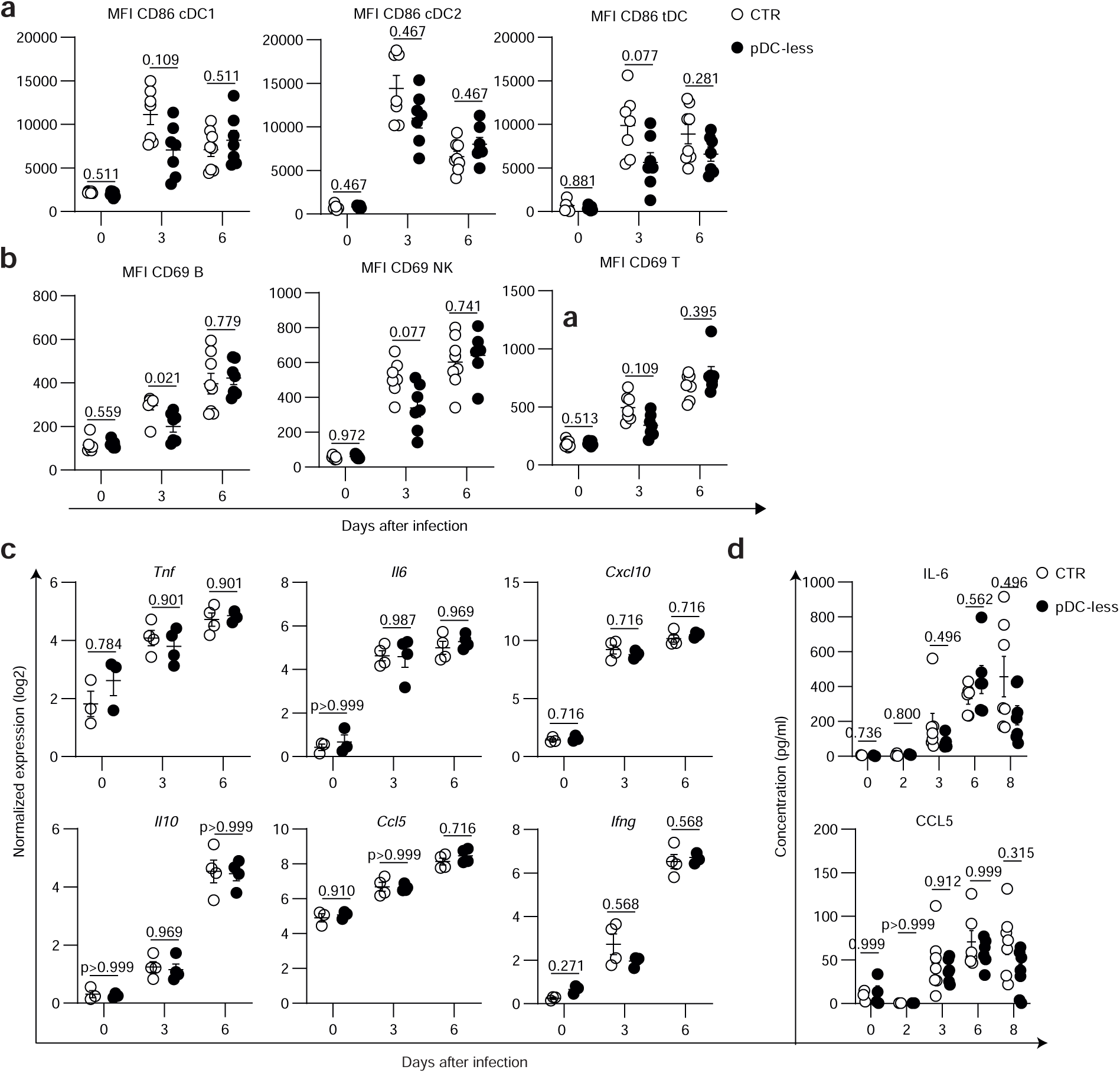
Activation state of immune cells and cytokine production in the lungs of control vs pDC-less mice during IAV infection. a-b, Mean Fluorescence intensity (MFI) of CD86 (**a**) and of CD69 (**b**) molecules expressed by indicated immune cell types isolated from the lungs of control or pDC-less mice at indicated days after IAV infection. **c**, Normalized expression (log2) of indicated cytokine genes based on Bulk RNASeq samples obtained by whole lung samples isolated from CTR and pDC-less mice at day 0 (n=3/strain), 3 and 6 (n=4/strain) after IAV infection. **d,** Cytometric bead arrays was performed to quantify cytokine concentrations in the BAL isolated from CTR and pDC-less mice uninfected (0) or at indicated days after infection. Each dot represents an individual animal. The data shown (mean +/- s.e.m.) are pooled from two independent experiments (n=3 CTR and 4 pDC-less at day 0, n=3 CTR and 2 pDC-less at day 2, n=7 at day 3, =6 at day 6, =7 for day 8 for both strains). An unpaired and nonparametric multiple t-test (Mann-Whitney) with Holm-Sidak method for correction was used for the statistical analysis.

**Extended Fig. 8:**
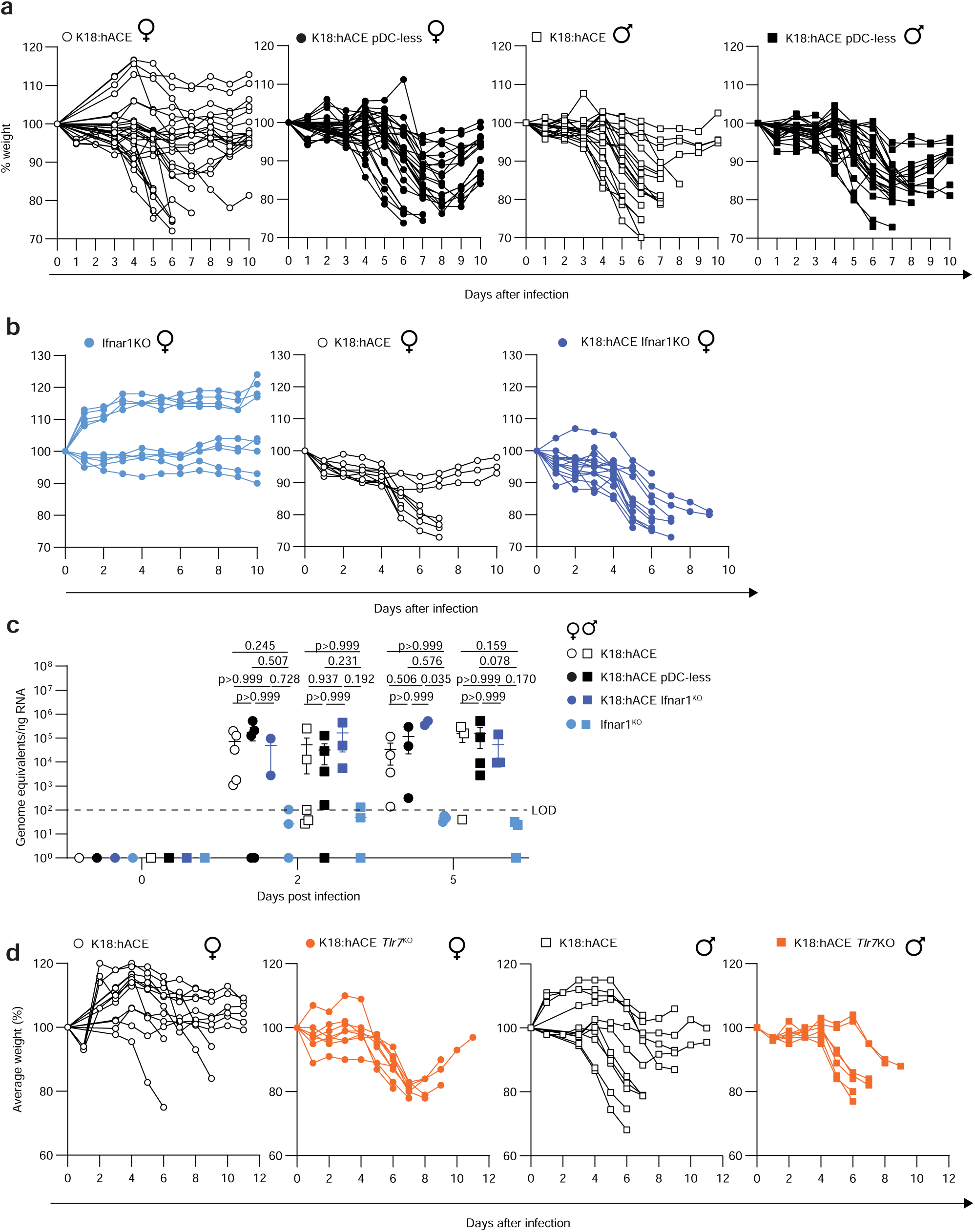
Impact of the loss of pDCs, *Ifnar1* or *Tlr7* on the weight curves of SARS-CoV2-infected K18:hACE. a-b, The weight of indicated SARS-CoV2-infected mice was regularly measured up to 12 days after infection. Data are represented as percentages of the weight at day 0 of the infection. Mice cohort numbers are indicated in the legend of Fig. 8a**-b**. **c,** SARS-CoV-2 viral burdens were measured by RT-qPCR as genome equivalent of RdRp protein/ng of total RNA of lungs isolated from K18:hACE mice at 0, 2 and 5 days post infection. Each plotted dot represents the viral burden for an individual animal and bars represent median +/- s.e.m. A nonparametric one-way ANOVA (Kruskal-Wallis test followed by Dunn’s post-hoc test) was used for the statistical analysis. **d**, The weight of indicated SARS-CoV2-infected mice was regularly measured up to 12 days after infection. Data are represented as percentages of the weight at day 0 of the infection. Mice cohort numbers are indicated in the legend of Fig. 8c.

**Extended Fig. 9:**
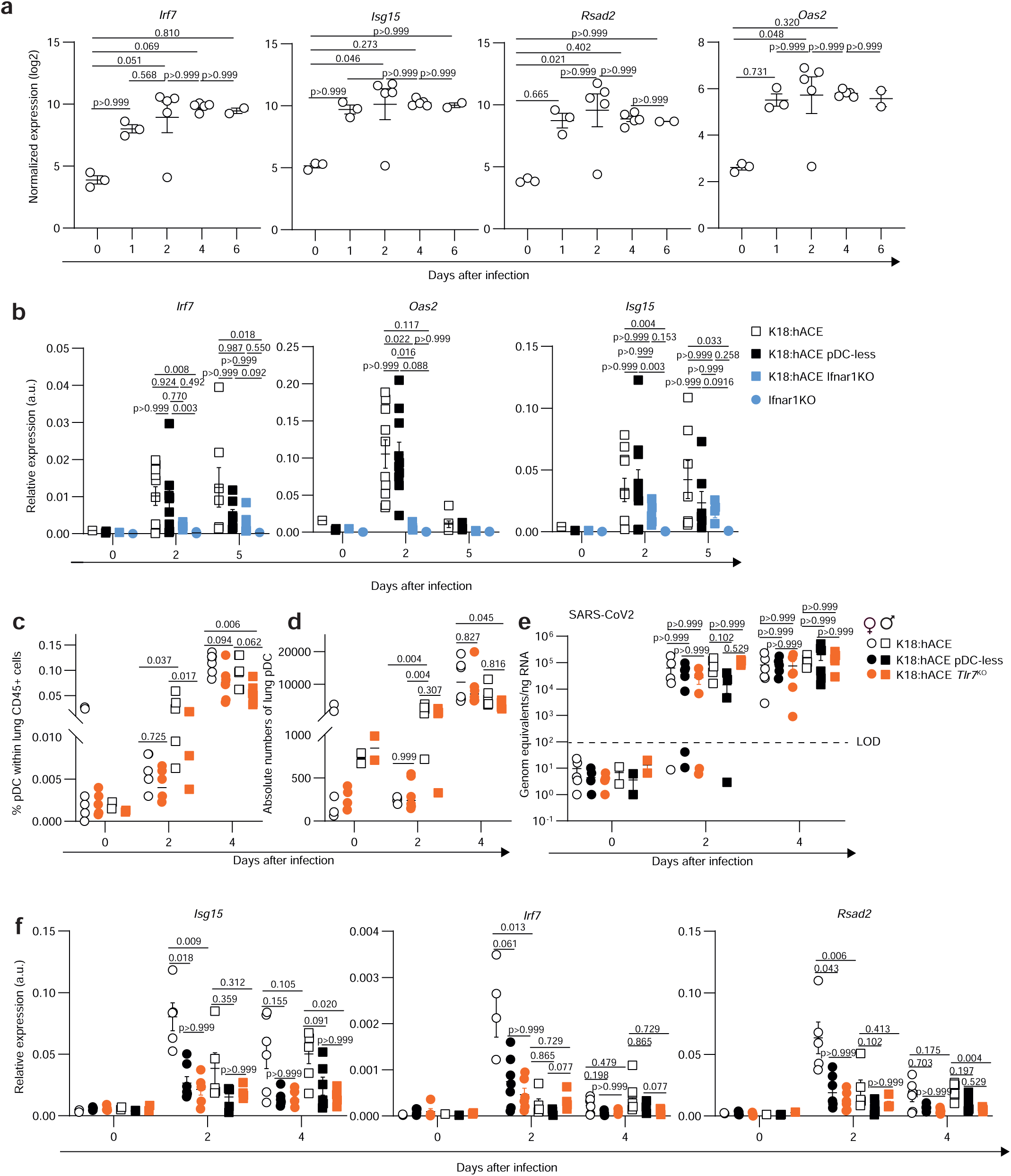
Interferon response and viral control regulation in SARS-CoV2-infected K18:hACE mice. **a**, Normalized expression (log2) of indicated genes based on Bulk RNASeq samples obtained by whole lung samples isolated from transgenic K18:hACE mice infected with 1.1 10^5^ PFU of SARS-CoV2 at indicated days after infection. Data (mean +/- s.e.m.) shown are from n= 3 at days 0 and 1, 5 at day 2 and 4, 2 at day 6 after infection. **b**, Expression levels of indicated ISGs were analyzed in indicated uninfected or SARS-CoV2-infected mice by RT-qPCR. The data (mean + s.e.m.) shown are from two independent experiments (n=2 for all strains at day 0, n=10 for K18:hACE and K18:hACE pDC-less, =5 for K18:hACE *Ifnar1*^KO^ and *Ifnar1*^KO^ at day 2, n= 7 for K18:hACE and K18:hACE pDC-less, =5 for K18:hACE *Ifnar1*^KO^ and =3 for *Ifnar1*^KO^ at day 5). **c-d**, % of pDCs within CD45^+^ lung cells (**c**) and absolute pDC numbers (**d**) in CTR and *Tlr7*^KO^ K18:hACE mice at indicated times after infection. **e,** SARS-CoV-2 viral burdens were measured by RT-qPCR as genome equivalent of RdRp protein/ng of total RNA of lungs isolated from K18:hACE mice at 0, 2 and 4 days post infection. Each plotted dot represents the viral burden for an individual animal and bars represent median +/- s.e.m. **f**, Expression levels of indicated genes was analyzed by RT-qPCR in lungs isolated from uninfected or SARS-CoV2-infected control (CTR), pDC-less and *Tlr7*^KO^ K18:hACE mice at indicated days after infection. Data were normalized to expression level of *Actin* gene. In **c-f**, at day 0 n= 7F and 2M CTR, 5F and 2M pDC-less, 4F and 2M *Tlr7*^KO^; at day 2, 5F and 5M CTR, 7F and 5M pDC-less, 6F and 5M *Tlr7*^KO^; at day 4, 6F and 6 M CTR for each strain). A nonparametric one-way ANOVA (Kruskal-Wallis test followed by Dunn’s post-hoc test) was used for the statistical analysis in **a-b, e-f** panels. A two-way ANOVA with Tukey’s multiple comparison test was used for the statistical analysis in **c-d** panels.

**Supplementary Table 1 :**
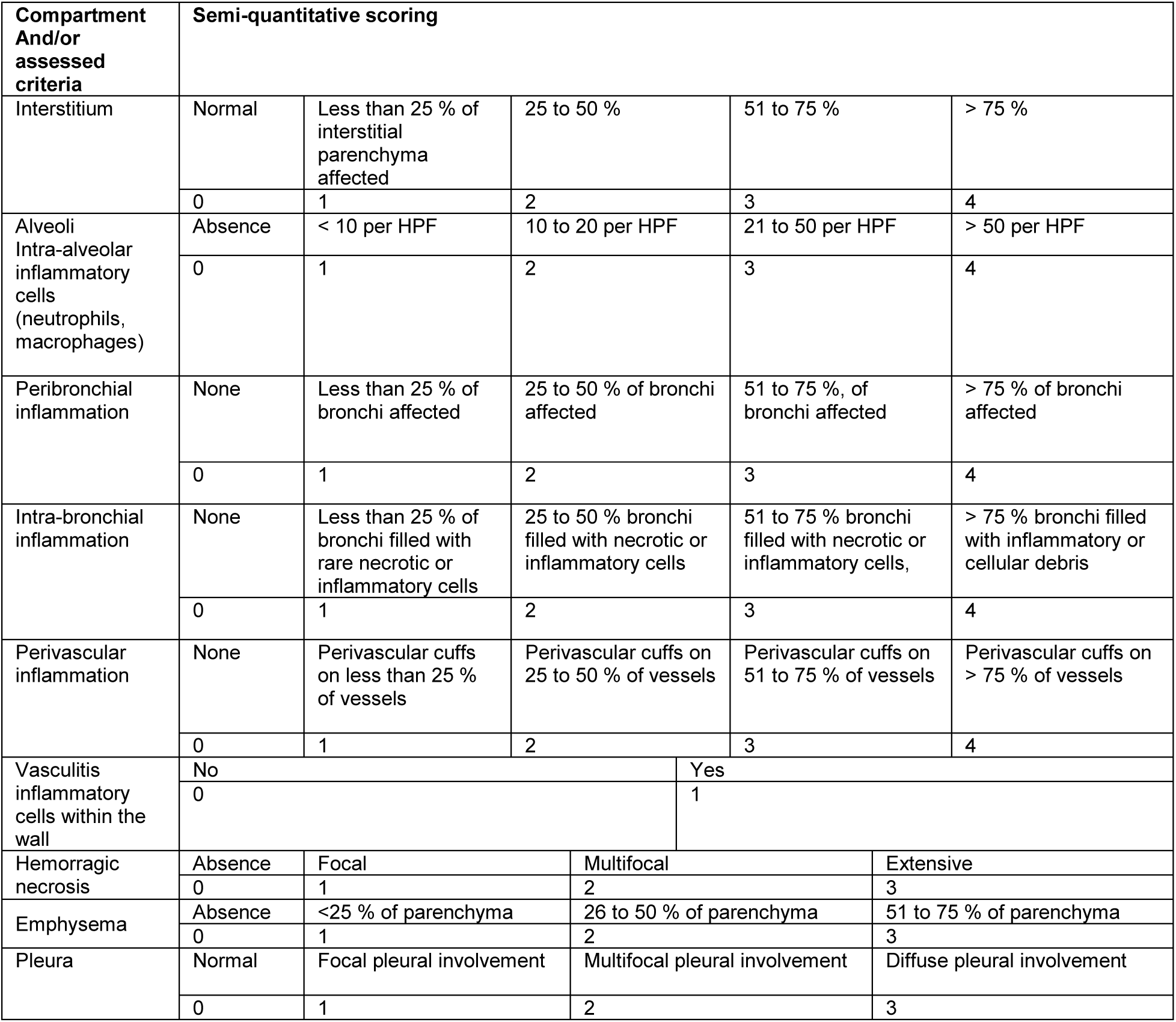
Semi-quantitative scoring of lung inflammation in IAV-infected mice.

## Supplementary methods

### RNA extraction and RT-qPCR on brain samples

Brains were harvested from euthanized mice after intracardiac perfusion with PBS 1x. Right hemispheres were then put in TRIzol (Invitrogen) and kept at -80°C until RNA extraction. Total RNAs cells were extracted following the protocol provided by the TRIzol Reagent user guide. The quantity of total RNA extracted was quantified using a Thermo Scientific NanoDrop. Purified RNA was reverse transcribed into cDNA by using an iScript cDNA Synthesis kit (Life Technologies).

The expression levels of the following murine genes: *Actin, Irf7, Isg15, Rsad2, S100a9, Ngp, Plp1, Dcx* as well as that of the influenza virus matrix protein gene (*M1*) were determined by quantitative PCR (qPCR) using the SYBR® Premix Ex TaqTM kit and analysed using the Prism 7500 Fast PCR System. Relative gene expression was calculated using the ΔΔCt method with *Actinb* as housekeeping gene for normalization. The primers used were as follows: *Actinb* forward 5’-GGCTGTATTCCCCTCCATCG-3’; reverse 5’-CCAGTTGGTAACAATGCCATGT-3’; *Irf7* forward 5’— CCACGCTATACCATCTACCTGG-3’ ; reverse 5’- GCTGCTATCCAGGGAAGACAC-3’ ; *Isg15* forward 5’— GGTGTCCGTGACTAACTCCAT-3’ ; reverse 5’-TGGAAAGGGTAAGACCGTCCT-3’ ; *Rsad2*forward 5’— TGCTGGCTGAGAATAGCATTAGG-3’ ; reverse 5’-GCTGAGTGCTGTTCCCATCT-3’ ; *M1 protein IAV* forward 5’-AAGACCAATCCTGTCACCTCTGA-3’; reverse 5’- CAAAGCGTCTACGCTGCAGTCC-3’ ; *S100a9* forward 5’-CAAATGGTGGAAGCACAGTT-3’ ; reverse 5’-AGCATCATACACTCCTCAAAGC-3’ ; *Ngp* forward 5’- AGACCTTTGTATTGGTGGTGGC-3’ ; reverse 5’-GGTTGTATGCCTCTATGGCTCTA-3’ ; *Plp1* forward 5’-TGGAGTCAGAGTGCCAAAGACA-3’ ; reverse 5’- GACATACTGGAAAGCATGAATCACA-3’ ; *Dcx* forward 5’-AACCAGAACCTTGCAGGCATTA- 3’ ; reverse 5’-CATAGCTTTCCCCTTCTTCCAGT-3’.

### Brain immunohistofluorescence, microscopy and image analysis

Mice were euthanized with overdose of ketamine-xylazine anesthetic and perfused transcardially with phosphate buffered saline (PBS) for antibody staining. Skull caps and meninges were removed, and following a brief wash in PBS, the brains were sectioned sagittally to keep left side for microscopic analysis and right side for RNA extraction. The left-brain parts were incubated for 2 days in formalin 2,5% diluted in PBS 1x (Sigma), then overnight in 30% sucrose solution diluted in PBS 1x (Sigma). Brains were then embedded in OCT freezing medium (CellPath), snap frozen and stored at −80 °C. 25 μm-thicked cryosections were performed using a microtome (Leica 3050s Cryostat) at temperatures between −23 °C and −20 °C.

For immunostaining, brain sections were blocked with PBS, 0.5% Triton X-100 and 2% Fetal Bovin Serum (FBS) for 1h at room temperature, then stained overnight at 4°C with primary antibodies diluted in PBS, 0.5% Triton X-100 and 2% FBS, with the exception of the anti-cFos antibody, which was incubated alone for 5h at 37°C before adding the other primary antibodies. The day after, brain sections were washed for 30 s and stained with secondary antibodies 2h at room temperature. Finally, brain sections were placed in mounting medium (FluorSave Reagent with DAPI) on a glass slide (Superfrost Plus, ThermoScientific) and cover-slipped.

Brain sections stained for anti-mouse NP, anti-mouse NeuN and anti-mouse cFos were acquired by confocal microscope (Zeiss LSM880 Indimo) with 10x. Brain sections stained for anti-mouse Iba1 and anti-mouse GFAP were acquired by slide scanner (Sysmex, Pannoramic Scan SC 150). Pictures were analyzed using QuPath.

**Legend Suppl. Figure 1:**
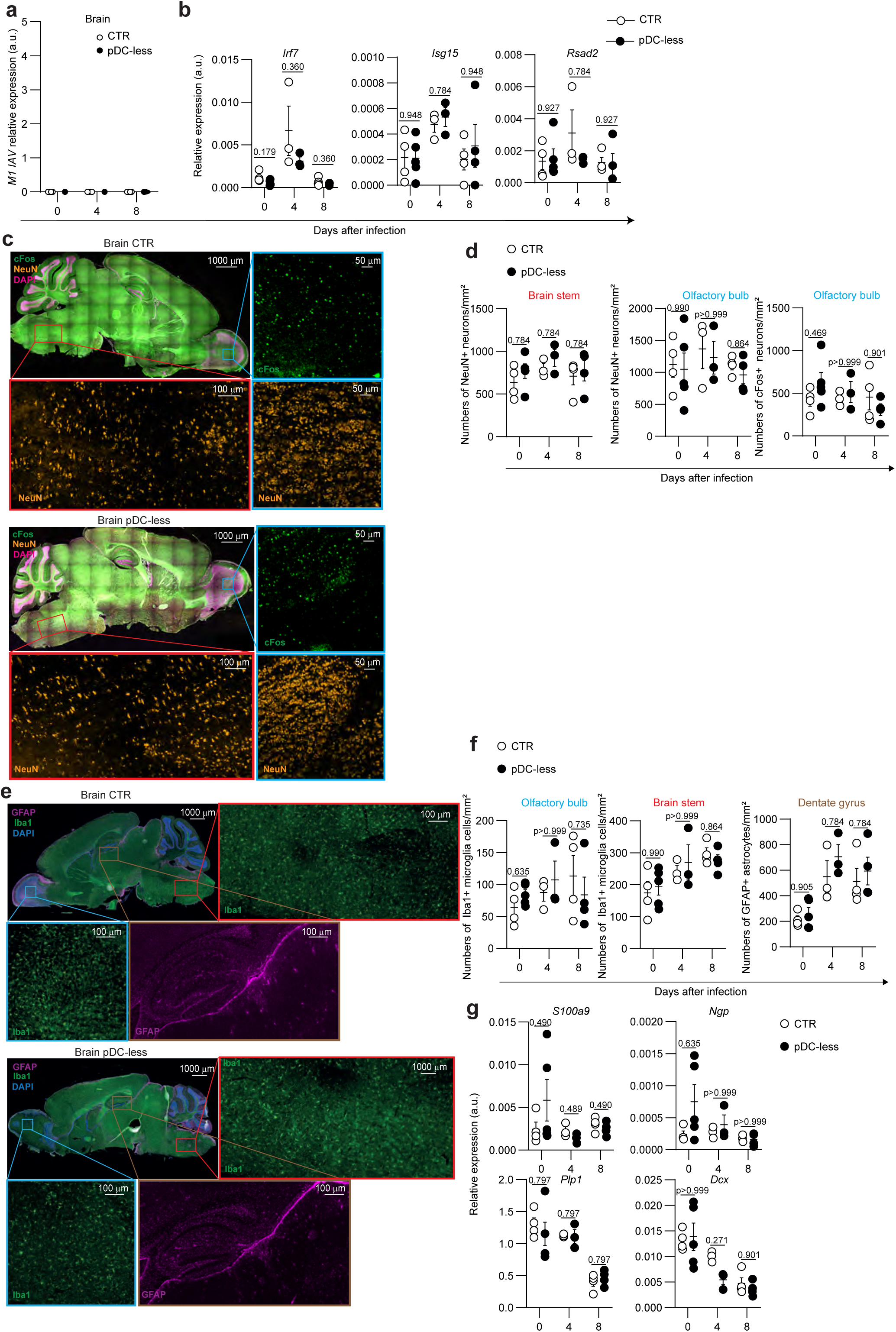
Histological and transcriptional analysis of brains isolated from uninfected versus IAV-infected control and pDC-less mice **a-b**, Expression levels of IAV *M1* gene (**a**) or of indicated ISGs (**b**) was analyzed by RT-qPCR in brains isolated from uninfected or IAV-infected CTR and pDC-less mice at indicated days after infection. Data were normalized to expression level of *Actin* gene. The data (mean + s.e.m.) shown are from two independent experiments (n=4 for each condition at each time points). **c-f**, Representative images of whole brains or of brain stem (red outline), olfactory bulb (blue outline) and dentate gyrus (brown outline) of indicated mice at day 4 after infection (**c,e**). **d, f**) Positive cells for indicated markers were quantified in indicated brain areas. First, the grid scale option was activated to determine the annotations of the following regions: brainstem (6x4 squares), olfactory bulb (2x2 squares) and dentate gyrus (5x3 squares) on the brain section. Then, for each region, the different cell types were quantified by cell detection with a threshold of 10 for NeuN, 10 for cFos, 12000 for GFAP and 10000 for Iba1. **g**, Expression levels of indicated genes was analyzed by RT-qPCR in brains isolated from uninfected or IAV-infected CTR and pDC-less mice at indicated days after infection. n=5 at day 0, 3 at day 4 and 4 at day 8 after infection.

## References

1. Tomasello, E., Pollet, E., Vu Manh, T.P., Uze, G. & Dalod, M. Harnessing Mechanistic Knowledge on Beneficial Versus Deleterious IFN-I Effects to Design Innovative Immunotherapies Targeting Cytokine Activity to Specific Cell Types. Front Immunol 5, 526 (2014).

2. Broggi, A. et al. Type III interferons disrupt the lung epithelial barrier upon viral recognition. Science 369, 706–712 (2020).

3. Major, J. et al. Type I and III interferons disrupt lung epithelial repair during recovery from viral infection. Science 369, 712–717 (2020).

4. Ngo, C., Garrec, C., Tomasello, E. & Dalod, M. The role of plasmacytoid dendritic cells (pDCs) in tissue immunity during viral infections and beyond. Cell Mol Immunol 21, 1008–1035 (2024).

5. Reizis, B. Plasmacytoid Dendritic Cells: Development, Regulation, and Function. Immunity 50, 37–50 (2019).

6. Bocharov, G. et al. A systems immunology approach to plasmacytoid dendritic cell function in cytopathic virus infections. PLoS Pathog 6, e1001017 (2010).

7. Reizis, B., Bunin, A., Ghosh, H.S., Lewis, K.L. & Sisirak, V. Plasmacytoid dendritic cells: recent progress and open questions. Annu Rev Immunol 29, 163–183 (2011).

8. Sa Ribero, M., Jouvenet, N., Dreux, M. & Nisole, S. Interplay between SARS-CoV-2 and the type I interferon response. PLoS Pathog 16, e1008737 (2020).

9. von Bernuth, H. et al. Pyogenic bacterial infections in humans with MyD88 deficiency. Science 321, 691–696 (2008).

10. Ku, C.L. et al. Selective predisposition to bacterial infections in IRAK-4-deficient children: IRAK-4-dependent TLRs are otherwise redundant in protective immunity. J Exp Med 204, 2407–2422 (2007).

11. Casanova, J.L. & Abel, L. Mechanisms of viral inflammation and disease in humans. Science 374, 1080–1086 (2021).

12. Ciancanelli, M.J. et al. Infectious disease. Life-threatening influenza and impaired interferon amplification in human IRF7 deficiency. Science 348, 448–453 (2015).

13. Zhang, Q. et al. Inborn errors of type I IFN immunity in patients with life-threatening COVID-19. Science 370 (2020).

14. Campbell, T.M. et al. Respiratory viral infections in otherwise healthy humans with inherited IRF7 deficiency. J Exp Med 219 (2022).

15. Zhang, Q. et al. Recessive inborn errors of type I IFN immunity in children with COVID-19 pneumonia. J Exp Med 219 (2022).

16. Asano, T., et al. X-linked recessive TLR7 deficiency in ∼1% of men under 60 years old with life-threatening COVID-19. Sci Immunol 6 (2021).

17. Garcia-Garcia, A. et al. Humans with inherited MyD88 and IRAK-4 deficiencies are predisposed to hypoxemic COVID-19 pneumonia. J Exp Med 220 (2023).

18. Mogensen, T.H. Human genetics of SARS-CoV-2 infection and critical COVID-19. Clin Microbiol Infect 28, 1417–1421 (2022).

19. Duncan, C.J.A. et al. Life-threatening viral disease in a novel form of autosomal recessive IFNAR2 deficiency in the Arctic. J Exp Med 219 (2022).

20. Cocita, C. et al. Natural Killer Cell Sensing of Infected Cells Compensates for MyD88 Deficiency but Not IFN-I Activity in Resistance to Mouse Cytomegalovirus. PLoS Pathog 11, e1004897 (2015).

21. Dalod, M. & Scheu, S. Dendritic cell functions in vivo: A user’s guide to current and next-generation mutant mouse models. Eur J Immunol 52, 1712–1749 (2022).

22. Allman, D. et al. Ikaros is required for plasmacytoid dendritic cell differentiation. Blood 108, 4025–4034 (2006).

23. Swiecki, M. et al. Cell depletion in mice that express diphtheria toxin receptor under the control of SiglecH encompasses more than plasmacytoid dendritic cells. J Immunol 192, 4409–4416 (2014).

24. Takagi, H. et al. Plasmacytoid dendritic cells are crucial for the initiation of inflammation and T cell immunity in vivo. Immunity 35, 958–971 (2011).

25. Cisse, B. et al. Transcription factor E2-2 is an essential and specific regulator of plasmacytoid dendritic cell development. Cell 135, 37–48 (2008).

26. Rodrigues, P.F. et al. Distinct progenitor lineages contribute to the heterogeneity of plasmacytoid dendritic cells. Nat Immunol 19, 711–722 (2018).

27. Leylek, R. et al. Integrated Cross-Species Analysis Identifies a Conserved Transitional Dendritic Cell Population. Cell Rep 29, 3736–3750 e3738 (2019).

28. Dress, R.J. et al. Plasmacytoid dendritic cells develop from Ly6D(+) lymphoid progenitors distinct from the myeloid lineage. Nat Immunol 20, 852–864 (2019).

29. Sulczewski, F.B. et al. Transitional dendritic cells are distinct from conventional DC2 precursors and mediate proinflammatory antiviral responses. Nat Immunol 24, 1265–1280 (2023).

30. Swiecki, M., Gilfillan, S., Vermi, W., Wang, Y. & Colonna, M. Plasmacytoid dendritic cell ablation impacts early interferon responses and antiviral NK and CD8(+) T cell accrual. Immunity 33, 955–966 (2010).

31. Stutte, S. et al. Type I interferon mediated induction of somatostatin leads to suppression of ghrelin and appetite thereby promoting viral immunity in mice. Brain Behav Immun 95, 429–443 (2021).

32. Valente, M. et al. Novel mouse models based on intersectional genetics to identify and characterize plasmacytoid dendritic cells. Nat Immunol 24, 714–728 (2023).

33. Le Goffic, R. et al. Detrimental contribution of the Toll-like receptor (TLR)3 to influenza A virus-induced acute pneumonia. PLoS Pathog 2, e53 (2006).

34. Bagadia, P. et al. An Nfil3-Zeb2-Id2 pathway imposes Irf8 enhancer switching during cDC1 development. Nat Immunol 20, 1174–1185 (2019).

35. Dalod, M. et al. Interferon alpha/beta and interleukin 12 responses to viral infections: pathways regulating dendritic cell cytokine expression in vivo. J Exp Med 195, 517–528 (2002).

36. Zucchini, N. et al. Individual plasmacytoid dendritic cells are major contributors to the production of multiple innate cytokines in an organ-specific manner during viral infection. Int Immunol 20, 45–56 (2008).

37. Abbas, A. et al. The activation trajectory of plasmacytoid dendritic cells in vivo during a viral infection. Nat Immunol 21, 983–997 (2020).

38. Dalod, M. et al. Dendritic cell responses to early murine cytomegalovirus infection: subset functional specialization and differential regulation by interferon alpha/beta. J Exp Med 197, 885–898 (2003).

39. Steinberg, C. et al. The IFN regulatory factor 7-dependent type I IFN response is not essential for early resistance against murine cytomegalovirus infection. Eur J Immunol 39, 1007–1018 (2009).

40. Baranek, T. et al. Differential responses of immune cells to type I interferon contribute to host resistance to viral infection. Cell Host Microbe 12, 571–584 (2012).

41. GeurtsvanKessel, C.H. et al. Clearance of influenza virus from the lung depends on migratory langerin+CD11b-but not plasmacytoid dendritic cells. J Exp Med 205, 1621–1634 (2008).

42. Wolf, A.I. et al. Plasmacytoid dendritic cells are dispensable during primary influenza virus infection. J Immunol 182, 871–879 (2009).

43. Kaminski, M.M., Ohnemus, A., Cornitescu, M. & Staeheli, P. Plasmacytoid dendritic cells and Toll-like receptor 7-dependent signalling promote efficient protection of mice against highly virulent influenza A virus. J Gen Virol 93, 555–559 (2012).

44. Davidson, S., Crotta, S., McCabe, T.M. & Wack, A. Pathogenic potential of interferon alphabeta in acute influenza infection. Nat Commun 5, 3864 (2014).

45. Davidson, S. et al. IFNlambda is a potent anti-influenza therapeutic without the inflammatory side effects of IFNalpha treatment. EMBO Mol Med 8, 1099–1112 (2016).

46. Rappe, J.C.F. et al. A TLR7 antagonist restricts interferon-dependent and -independent immunopathology in a mouse model of severe influenza. J Exp Med 218 (2021).

47. Scheu, S., Dresing, P. & Locksley, R.M. Visualization of IFNbeta production by plasmacytoid versus conventional dendritic cells under specific stimulation conditions in vivo. Proc Natl Acad Sci U S A 105, 20416–20421 (2008).

48. Tomasello, E. et al. Molecular dissection of plasmacytoid dendritic cell activation in vivo during a viral infection. EMBO J 37 (2018).

49. Creusat, F. et al. IFN-γ primes bone marrow neutrophils to acquire regulatory functions in severe viral respiratory infections. Sci Adv 10, eadn3257 (2024).

50. Chen, H. et al. Pulmonary permeability assessed by fluorescent-labeled dextran instilled intranasally into mice with LPS-induced acute lung injury. PLoS One 9, e101925 (2014).

51. Lucas, C. et al. Longitudinal analyses reveal immunological misfiring in severe COVID-19. Nature 584, 463–469 (2020).

52. Blanco-Melo, D. et al. Imbalanced Host Response to SARS-CoV-2 Drives Development of COVID-19. Cell 181, 1036–1045 e1039 (2020).

53. Lee, J.S., et al. Immunophenotyping of COVID-19 and influenza highlights the role of type I interferons in development of severe COVID-19. Sci Immunol 5 (2020).

54. Laurent, P., et al. Sensing of SARS-CoV-2 by pDCs and their subsequent production of IFN-I contribute to macrophage-induced cytokine storm during COVID-19. Sci Immunol 7, eadd4906 (2022).

55. McCray, P.B., Jr., et al. Lethal infection of K18-hACE2 mice infected with severe acute respiratory syndrome coronavirus. J Virol 81, 813–821 (2007).

56. Takahashi, T. et al. Sex differences in immune responses that underlie COVID-19 disease outcomes. Nature 588, 315–320 (2020).

57. Channappanavar, R. et al. IFN-I response timing relative to virus replication determines MERS coronavirus infection outcomes. J Clin Invest 129, 3625–3639 (2019).

58. Channappanavar, R. et al. Dysregulated Type I Interferon and Inflammatory Monocyte-Macrophage Responses Cause Lethal Pneumonia in SARS-CoV-Infected Mice. Cell Host Microbe 19, 181–193 (2016).

59. Ullah, T.R. et al. Pharmacological inhibition of TBK1/IKKepsilon blunts immunopathology in a murine model of SARS-CoV-2 infection. Nat Commun 14, 5666 (2023).

60. Vu Manh, T.P., et al. Defining Mononuclear Phagocyte Subset Homology Across Several Distant Warm-Blooded Vertebrates Through Comparative Transcriptomics. Front Immunol 6, 299 (2015).

61. Swiecki, M., Wang, Y., Gilfillan, S. & Colonna, M. Plasmacytoid dendritic cells contribute to systemic but not local antiviral responses to HSV infections. PLoS Pathog 9, e1003728 (2013).

62. Jamali, A. et al. Characterization of Resident Corneal Plasmacytoid Dendritic Cells and Their Pivotal Role in Herpes Simplex Keratitis. Cell Rep 32, 108099 (2020).

63. Brewitz, A. et al. CD8(+) T Cells Orchestrate pDC-XCR1(+) Dendritic Cell Spatial and Functional Cooperativity to Optimize Priming. Immunity 46, 205–219 (2017).

64. Onodi, F. et al. SARS-CoV-2 induces human plasmacytoid predendritic cell diversification via UNC93B and IRAK4. J Exp Med 218 (2021).

65. Hadjadj, J. et al. Impaired type I interferon activity and inflammatory responses in severe COVID-19 patients. Science 369, 718–724 (2020).

66. Venet, M. et al. Severe COVID-19 patients have impaired plasmacytoid dendritic cell-mediated control of SARS-CoV-2. Nat Commun 14, 694 (2023).

67. Greene, T.T. & Zuniga, E.I. Type I Interferon Induction and Exhaustion during Viral Infection: Plasmacytoid Dendritic Cells and Emerging COVID-19 Findings. Viruses 13 (2021).

68. Bruel, T. et al. Plasmacytoid dendritic cell dynamics tune interferon-alfa production in SIV-infected cynomolgus macaques. PLoS Pathog 10, e1003915 (2014).

69. Mitchell, J.L. et al. Plasmacytoid dendritic cells sense HIV replication before detectable viremia following treatment interruption. J Clin Invest 130, 2845–2858 (2020).

70. Buitendijk, M., Eszterhas, S.K. & Howell, A.L. Gardiquimod: a Toll-like receptor-7 agonist that inhibits HIV type 1 infection of human macrophages and activated T cells. AIDS Res Hum Retroviruses 29, 907–918 (2013).

71. Meng, F.Z. et al. TLR7 Activation of Macrophages by Imiquimod Inhibits HIV Infection through Modulation of Viral Entry Cellular Factors. Biology (Basel) 10 (2021).

72. Marie, I., Durbin, J.E. & Levy, D.E. Differential viral induction of distinct interferon-alpha genes by positive feedback through interferon regulatory factor-7. EMBO J 17, 6660–6669 (1998).

73. Prakash, A. & Levy, D.E. Regulation of IRF7 through cell type-specific protein stability. Biochem Biophys Res Commun 342, 50–56 (2006).

74. Yang, K. et al. Human TLR-7-, -8-, and -9-mediated induction of IFN-alpha/beta and - lambda Is IRAK-4 dependent and redundant for protective immunity to viruses. Immunity 23, 465–478 (2005).

75. Zhang, Q. et al. Autoantibodies against type I IFNs in patients with critical influenza pneumonia. J Exp Med 219 (2022).

76. Piersma, S.J. et al. Virus infection is controlled by hematopoietic and stromal cell sensing of murine cytomegalovirus through STING. Elife 9 (2020).

77. Kadoki, M. et al. Organism-Level Analysis of Vaccination Reveals Networks of Protection across Tissues. Cell 171, 398–413 e321 (2017).

78. Beura, L.K. et al. Normalizing the environment recapitulates adult human immune traits in laboratory mice. Nature 532, 512–516 (2016).

79. Reese, T.A. et al. Sequential Infection with Common Pathogens Promotes Human-like Immune Gene Expression and Altered Vaccine Response. Cell Host Microbe 19, 713–719 (2016).

80. Schneider, C.A., Rasband, W.S. & Eliceiri, K.W. NIH Image to ImageJ: 25 years of image analysis. Nat Methods 9, 671–675 (2012).

81. Bankhead, P. et al. QuPath: Open source software for digital pathology image analysis. Sci Rep 7, 16878 (2017).

